# Perceptual processes as charting operators

**DOI:** 10.1101/2025.06.25.661475

**Authors:** Peter Neri

## Abstract

Sensory operators are classically modelled using small circuits involving canonical computations, such as energy extraction and gain control. Notwithstanding their utility, circuit models do not provide a unified framework encompassing the variety of effects observed experimentally. We develop a novel, alternative framework that recasts sensory operators in the language of intrinsic geometry. We start from a plausible representation of perceptual processes that is akin to measuring distances over a sensory manifold. We show that this representation is sufficiently expressive to capture a wide range of empirical effects associated with elementary sensory computations. The resulting geometrical framework offers a new perspective on state-of-the-art empirical descriptors of sensory behavior, such as first-order and second-order perceptual kernels. For example, it relates these descriptors to notions of flatness and curvature in perceptual space.

## 1 Introduction

In order for quantitative accounts of perception to be practically viable, relevant empirical characterizations must be restricted to settings in which animals respond to sensory stimuli by producing measurable motor outputs. In this study, we further restrict our characterization to the simplified scenario in which there are only two possible classes of stimuli, such as “predator” or “prey”, and two motor outputs, such as “fight” or “flight”. We will also restrict our characterization to conditions under which the choice between these two alternatives is only driven by the nature of the input stimulus (i.e. it does not depend, for example, on whether the animal is hungry or thirsty). Under this simplified scenario, we can describe the underlying process in the form of a functional H that maps input vector **s** (stimulus) onto binary response *r* (motor output): *r*= *ℋ* (**s**).

There are many different ways in which we can study this process experimentally. However, if we are to provide a quantitative characterization of its properties, there is essentially only one way to go about it: psychophysics (Green and Swets, 1966). In psychophysical experiments, different instances of **s** are presented to the animal, and each instance is associated with a binary output *r*. By collecting many such instances of stimulus-response associations, it is hoped that enough information can be gathered to constrain a representation of *ℋ* that adequately and parsimoniously predicts *r* in response to arbitrary **s**. The term “adequately” means: to saturation of the boundary imposed on prediction by stimulus-independent variability that is intrinsic to the animal (Faisal et al., 2008; Neri, 2010a).

*ℋ* can be represented using a diverse range of frameworks and tools (Barack and Krakauer, 2021; Langdon et al., 2023). For example, one may use analytical expressions incorporating a summary characteristic of the stimulus (Zhou et al., 2024) (e.g. contrast), or one may specify a biologically plausible circuit that operates directly on the input image (Schütt and Wichmann, 2017; Lyu et al., 2021). More recently, a common strategy has been to learn the stimulus-response mapping via neural networks that, in some cases, spontaneously acquire properties resembling those associated with specific biological structures (Yamins and DiCarlo, 2016; Neri, 2022). In this article, we adopt an altogether different approach that relies on geometric constructs (Amari, 2016; Petitot, 2017; Chung and Abbott, 2021; Neri, 2024). Although not transparently relatable to physiological structures, this approach offers a novel unified perspective that is able to account for various empirical observations. Furthermore, this perspective carries the potential to seamlessly integrate those observations into a wider-scale account of sensory computation encompassing a much larger class of perceptual phenomena.

## 2 Theoretical framework

### 2.1 From stimulus set to Riemannian manifold structure

The most fundamental assumption underlying our theoretical framework is that the set of all possible stimuli that can be generated for a given experiment can be equipped with Riemannian manifold structure. To establish a topology on this set, we inherit the standard topology from the real-valued vector space associated with the manner in which we store and manipulate stimuli on the computer. We therefore start by formalizing how the latter process is structured in the laboratory.

When we operate on the computer for the purpose of generating stimuli and analyzing them, each stimulus is associated with an element in ℝ^*n*^ (up to the precision of our rendering device), and we can perform addition and multiplication on a collection of such elements. The resulting vector space is not closed under those two operations: a physically realizable generator cannot produce any arbitrary stimulus due to hardware limitations, however we assume that vector space structure applies to a satisfactory degree of approximation within the operating range of the experiments. In other words, we assume judicious specification of stimulus parameterization such that hardware capability is not exceeded.

The atlas (collection of charts) is established by the perceptual process, and so is a metric structure with an inner product. When perception is queried for the purpose of obtaining a perceptal judgment, it selects a specific chart, which maps the input stimulus to a collection of real coordinate values, and performs measurements on said chart, which are then used to generate a behavioral decision. To simplify notation, we start by adopting chart coordinates that are trivially aligned with the stimulus values stored on the computer (Section 2.2), however our framework can accommodate other choices of chart coordinates (we discuss this issue further in Section 5.4).

We make one last assumption: the human observer stores templates of stimuli to-be-discriminated in memory. For the specific examples considered in this study, one template is associated with a noiseless stimulus containing the target signal to-be-detected, while the other template is associated with a noiseless stimulus *not* containing the target signal (detection task). More generally, the latter may be associated with a noiseless stimulus containing some non-target signal to be discriminated from the target signal (discrimination tasks, not included among the examples considered in this study). These templates map to two elements of the manifold, which essentially amounts to stating that the perceptual templates stored in memory by the perceptual process can be realized, at least in principle, on stimulus hardware.

### 2.2 Stimulus/template specification with associated notation

With the above framework in mind, each stimulus vector **s** is specified by a collection of real coordinate values *s*^*i*^ on the chart instantiated by the perceptual process. For the examples considered in this study, the stimulus is constructed by combining two component vectors: a signal component and a noise component (see **Figure 1D** for a specific example). The signal component can only take one of two pre-specified configurations: the “target-present” configuration **t**, and the “target-absent” configuration 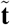. In a typical detection task such as those considered here, **t** may correspond to a bright bar in the middle of the monitor (**Figure 1C** shows an example embedded in visual noise), while 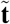 may correspond to the featureless background. The above (and further detailed below) stimulus specifications are not necessary for the framework outlined in Sections 2.3 and 2.4 to hold; for example, stimuli may not contain noise, and/or may be characterized by natural statistics. For the purpose of kernel derivation, however, the classic signal-in-white-noise approach (Marmarelis and Marmarelis, 1978; Schetzen, 1980) is highly preferable because it is amenable to analytical treatment (Sections 2.5 and 2.6), and it retains a greater degree of interpretability (Sharpee et al., 2008; Landolfi and Neri, 2025).

**Figure 1:**
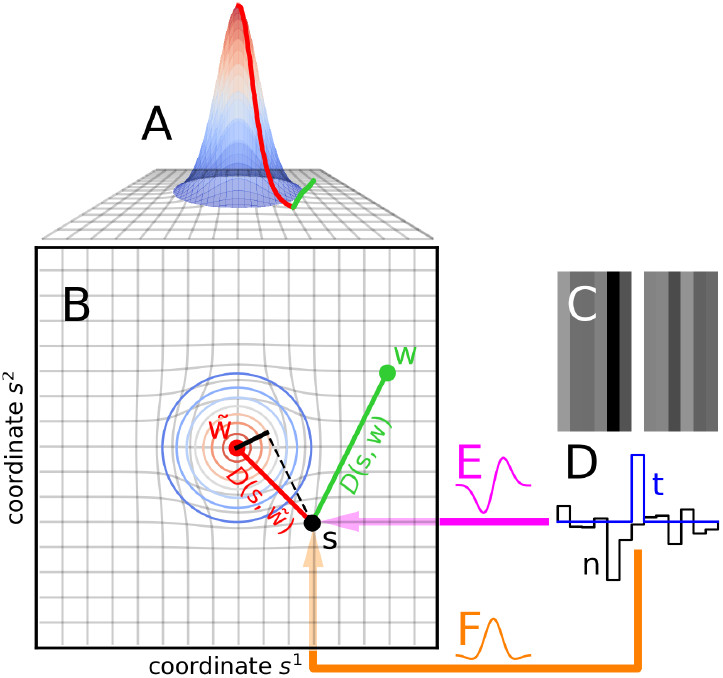
Stimulus **s** maps to a vector (black dot) in metric perceptual space (**B**). Memory templates map to vectors **w** (green dot) for target-to-be-detected, and 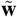 (red dot) for absence of said target. The perceptual system measures distance *D*(**s, w**) between stimulus **s** and template **w** (green segment), and distance *D*(**s**, 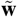) between **s** and 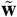 (red segment). Based on these measurements, the system can establish whether **s** is closer to **w** or to 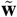, and classify it accordingly. In flat space (**B**), taking the difference between *D*(**s, w**) and *D*(**s**, 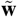) is equivalent (i.e. returns same perceptual decision) to measuring the projection of **s** onto **w** (black segment). For the example shown here, **s** is closer to 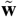 than it is to **w** (red segment is shorter than green segment). In curved space (**A**), distance measurements can lead to stimulus classifications that differ from those obtained in flat space: **s** is now closer to **w** than it is to 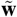 (green path is shorter than red path). To visualize higher-dimensional stimuli (**C**) on the plane (**B**), we project them onto vectors **E,F**. Each stimulus consists of two components: target and noise (**t,n** respectively in **D**).

The noise component **n** is different for every instance of **s**. More specifically, each noise coordinate *n*^*i*^ is independently drawn from a zero-mean Gaussian distribution. Locally, each stimulus coordinate is specified by either *s*^*i*^=*t*^*i*^+*n*^*i*^ when the stimulus is presented in target-present configuration, or 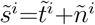 when the stimulus is presented in target-absent configuration. For the data and simulations presented in this study, every component of 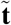 is (without loss of generality) specified to be zero (hence the term “target-absent”): 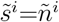. Our considerations, however, apply to arbitrary specifications of 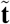 (discrimination tasks), so that the term target-absent may in fact be adopted with reference to the presence of a non-target signal (as anticipated in Section 2.3).

The two memory templates stored by the human observer, one for the target-present configuration and another one for the target-absent configuration, are denoted by vectors **w** and 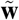, respectively. In general, **w**≠**t** and 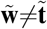: the memorized representations of target-present and target-absent stimuli, which exist in the head of the human observer, are not necessarily aligned with the physical specifications of those stimuli, which exist on the hardware that is used to display them.

We draw attention to a slight abuse of the 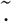 notation above. It is always true that 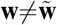 (the system otherwise carries no discriminative power). In general, it is also the case that 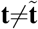 (except when specified otherwise by the experimenter, see for example Section 3.2 and **Figure 2C,F**). However, ⟨**n**⟩ = ⟨**ñ**⟩ where ⟨.⟩ is used to denote average over many trials (and = is intended asymptotically). In other words, the statistical specification of the noise component **n** is identical between target-present and target-absent stimuli: when applied to the noise component, the 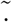 notation is used to refer to noise samples **ñ** associated with the target-absent stimulus 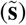 to distinguish them from those (**n**) associated with the target-present stimulus (**s**), however these two different types of noise samples conform to identical statistical specifications. In particular, ⟨*n*^*i*^⟩ = ⟨*ñ*^*i*^ ⟩=0 (zero-mean), ⟨*n*^*i*^*n*^*j*^ ⟩ = ⟨*ñ*^*i*^*ñ*^*j*^⟩ =*δ*_*ij*_ (uncorrelated coordinate values), and ⟨*n*^*i*^*ñ*^*j*^ ⟩ =0 (uncorrelated samples).

**Figure 2:**
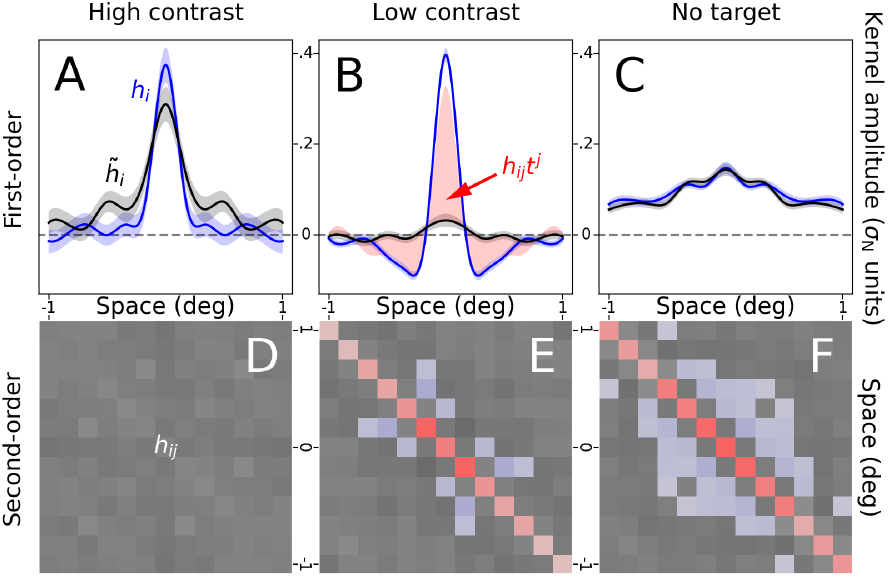
Empirically measured first-order (top row) and second-order (bottom row) kernels may vary independently as a function of specific stimulus manipulations. At high stimulus contrast (left column), target-present (blue) and target-absent (black) first-order kernels are not experimentally distinguishable (**A**), while the second-order kernel is featureless (**D**). At low contrast (middle column), the opposite pattern is observed: target-present and target-absent kernels are markedly different (**B**), while the second-order kernel is highly structured (**E**). When the target signal is removed from the stimulus (right column), target-present and target-absent kernels become indistinguishable by definition (and experimentally, see **C**), however the second-order kernel remains highly structured (**F**). Shaded regions in **A**–**C** show ± 1 SEM. Red/blue-tinted pixels in **D**–**F** indicate significant positive/negative modulations (|Z-score| *>*2). Red-tinted area in **B** shows theoretically predicted difference between blue and black traces (equation 23, Section 3.5). Kernel amplitude in **A**–**C** (y axis) is plotted in units of external noise SD. Each of the first two columns is computed from ≈ 35K trials of published data (Neri, 2018b) aggregated across 8 observers. Third column is computed from ≈ 32K trials of unpublished data aggregated across 10 observers.

### 2.3 Perceptual distance

Perceptual judgments are based on distance measurements on the stimulus manifold. To perform such measurements, we posit the existence of a metric with associated inner product. For a given choice of chart and at a given point on the manifold, we can express this product as *g*_*ij*_*x*^*i*^*y*^*j*^ (Einstein notation), where *x*^*i*^/*y*^*i*^ are the chart coordinates of vectors **x/y**. Using the inner product, we can readily define kinetic energy along a curve *γ* running between stimulus **s** and memory template **w** over the manifold. With *γ*(0)=**w** and *γ*(1)=**s**, this quantity can be expressed as:

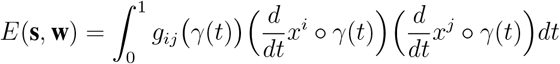

We assume that the perceptual process can measure this quantity along a geodesic path to obtain an energy-based measure of distance *D*:

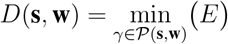

where *𝒫* (**s, w**) is the collection of all curves running between **s** and **w**. The issue of whether and how the perceptual process may be able to identify a geodesic path is complex, and we return to this point in Section 5.4.

Without detailed knowledge of the metric specification (which we cannot acquire from isolated behavioral experiments, see Section 5.1), the above expressions are useless for applied purposes. To take a first step in the direction of deriving useful expressions (to be further developed in Section 2.6), we notice that, because *D* is a scalar value, we can always specify a matrix *g* such that the same value is returned by the simpler expression:

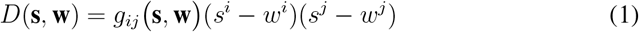

where *g*_*ij*_(**s, w**) can, in principle, differ for every choice of **s** and **w**. We refer to *g* as the “ambient metric”, because *g* no longer corresponds to the metric at a given point on the manifold, but rather summarizes metric structure along a geodesic path between **s** and **w**.

The perceptual process only computes two types of distance *D*: between stimulus **s** (or 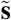) and target-present template **w**, and between **s** (or 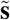) and target-absent template 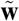. We reserve the symbol *g* for the former, and 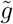 for the latter. We can therefore write *D*(**s, w**)=*g*_*ij*_(**s**)(*s*^*i*^ − *w*^*i*^)(*s*^*j*^ − *w*^*j*^) with the dependence of *g* on **w** implicit, and similarly 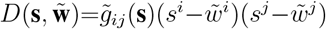 with the dependence of 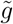 on 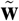 implicit.

To visualize the above procedure, we imagine that the perceptual system “places itself” at the element corresponding to **s** on the manifold (black point in **Figure 1B**). From this vantage point, it traces out a geodesic path to **w** (green path/segment in **Figure 1A/B**) along a Cartesian coordinate chart (i.e. a chart aligned with the geodesic), and measures *D*(**s, w**) using metric components that are collectively summarized by *g*_*ij*_. It then repeats this procedure with respect to 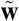 (red path/segment in **Figure 1A/B**) to obtain *D*(**s**, 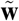) using metric components summarized by 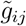. Clearly, it is meaningless to speak of the perceptual system in the manner verbalized above (i.e. as a homunculus operating within itself). This description is adopted here for intuitive purposes only, with no implication of any resemblance to the natural phenomenon.

### 2.4 Decision variables

We restrict our analysis to the following experimental scenario (two alternative forced choice protocol (Green and Swets, 1966)): on each trial, the human observer is presented with both **s** and 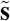, and is asked to select **s**. In the mind of the human observer, selecting **s** must be understood as selecting, between **s** and 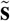, the stimulus most closely resembling **w** and least closely resembling 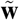.

In line with established accounts of perceptual decision-making (Pritchett and Murray, 2015), we posit that the human observer assigns a scalar value to each stimulus reflecting the likelihood that said stimulus is **s**. We refer to this value with the term “decision variable”. The decision variable associated with the target-present stimulus is *R*(**s**), while the decision variable associated with the target-absent stimulus is 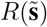. The observer selects the stimulus associated with largest decision variable as corresponding to **s**. Therefore, if 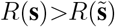 the observer will respond correctly, otherwise he/she will respond incorrectly.

Because the human observer is intrinsically noisy (Neri, 2010a), we must model human decision-making probabilistically (Green and Swets, 1966; Neri, 2010b; Carandini, 2024). We write the probability that a given response is correct as:

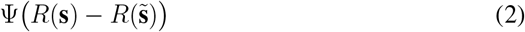

where we assume that the static nonlinear transducer Ψ only depends on the difference between *R*(**s**) and 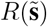 (Neri, 2013; Pritchett and Murray, 2015). A typical choice for Ψ is the cumulative Gaussian distribution function (Green and Swets, 1966), however there is empirical evidence that other distributions may capture perceptual noise more accurately (Neri, 2013; Sanborn et al., 2025) (see also Section A.2).

We consider three rules for generating the decision variable *R*. The first rule reflects the most commonly adopted model in computational neuroscience (Heeger et al., 1996; Dayan and Abbott, 2001; Ostojic and Brunel, 2011): the linear-nonlinear (LN) cascade, also termed template-matching in the behavioral literature (Brunelli and Poggio, 1997). The corresponding decision variable is:

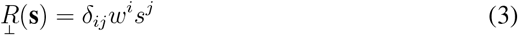

The above operation, which amounts to projecting stimulus **s** onto the target-present template **w** stored in memory by the observer (hence symbol ⊥ standing for “projection rule”), is entirely linear, however it is (inevitably) followed by the nonlinear operator Ψ (expression 2). This rule does not refer back to the metric *g*, in fact it is not even sensibly defined in the context of the geometrical framework we introduced earlier, except for the specific case of a flat geometry, for which *g* does not vary across the manifold. In that case, we could write 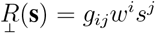, however this is equivalent to equation 3 with *g* absorbed into **w**.

The second rule generalizes the “similarity” rule proposed and studied by some previous authors (Watson et al., 1997; Solomon, 2002), and reflects the naive strategy of simply measuring the distance between stimulus **s** and **w**. The corresponding decision variable is:

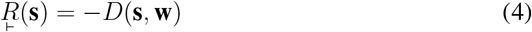

where ⊢ stands for “absolute distance rule”. The minus sign is introduced because the observer must now select the stimulus associated with smallest, not largest, *D*(**s, w**) (the smaller this quantity, the closer the stimulus to the target-present template).

The third rule is similar to ⊢ above, except it explicitly takes into account the memory template 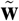 associated with the target-absent signal. This element is often neglected in perceptual models of detection tasks, because the target-absent stimulus typically consists of a blank screen, and is therefore regarded as a non-stimulus that has no impact on the final outcome (but see Beard and Ahumada (1999), in particular their Figure 5, for a decisional rule that explicitly incorporates the target-absent memory template). For example, if 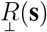 were reformulated to incorporate the term 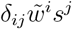, this term would have no impact because equal to zero for 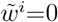. In the present framework, 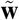 can always be set to 0 by redefining **w** as 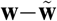. However, particularly when observers engage in two-way classification judgments, it is reasonable to posit that the stimulus-to-be-classified should be compared against both target-present and target-absent memory templates, to adjucate whether it is more similar to the former or to the latter. We can formulate the corresponding decision variable as:

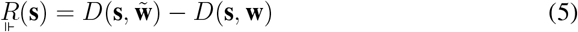

where ⊩ stands for “relative distance rule”. Notice that, in the expression above, the term *D*(**s**, 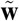) does not necessarily equal zero if we choose chart coordinates centred on 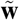. This is the essential characteristic that sets 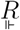 apart from 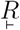: these two rules cannot be rendered equivalent by simply shifting the coordinate frame.

### 2.5 Kernel computation from data

To characterize *ℋ* in the laboratory, we must derive empirical descriptors of its characteristics. We focus on two established classes of descriptors (Murray, 2011): first-order (Ahumada, 2002) and second-order kernels (Neri, 2010b). These descriptors are not arbitrary: they represent the only logical approach for providing a transparent statistical description of input-output coupling in elementary perceptual judgments (Murray, 2011; Neri, 2018a). In this sense, we are not focusing here on some exotic, abstruse empirical representation of questionable relevance. To the contrary, we are taking as point of reference an indispensable class of descriptors for characterizing noise-based measurements of perceptual processing (Ahumada, 2002; Murray, 2011; Neri, 2018a).

For the first-order class, we compute target-present component *h*_*i*_ and target-absent component 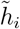 separately (Ahumada and Lovell, 1971) (see Section 3.1 for theoretical and empirical motivations behind this distinction). For the second-order class, we only compute the composite descriptor *h*_*ij*_ incorporating both target-present and target-absent components (Neri, 2004) (see Section 2.6 for theoretical justification of this latter choice).

We derive first-order kernels via the following calculations (Ahumada, 2002):

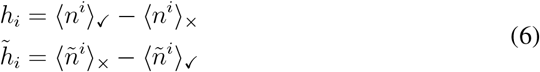

where subscript ✓ (or ×) refers to the subset of trials on which the observer responded correctly (or incorrectly).

To compute the second-order kernel, we replace averaging ⟨*n*^*i*^⟩ with covariance ⟨*n*^*i*^*n*^*j*^⟩−⟨*n*^*i*^⟩⟨*n*^*j*^⟩ in the two expressions above (Neri, 2004), and sum them to obtain:

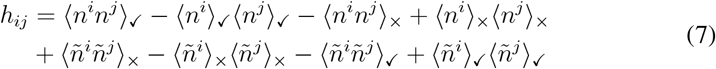

These empirical descriptors share some characteristics with their electrophysiological counterparts, the spike-triggered average (STA) (Rieke, 1999) and spike-triggered covariance (STC) (Schwartz et al., 2002), however they also present important differences. In particular, the second-order kernel above is characterized by a difference operation in which covariance matrices are subtracted for some response classes (*h*_*ij*_ is not necessarily expected to be positive definite, or semi-definite for correlated inputs, while STC always is). This simple fact has fundamental implications for its geometrical interpretation (Section 2.7) that set it apart from STC (see also Sandler and Marmarelis (2015)). In the case of first-order kernels, the difference operation is also present (trailing terms in equations 6), however it bears little conceptual significance for the comparison between these kernels and the STA.

### 2.6 Kernel derivation under plausible approximations

Because the decisional rules considered in Section 2.4 only contain quadratic elements, their differential output 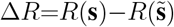 can be written as:

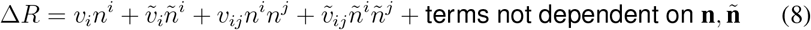

which is a simplified second-order Volterra expansion in **n** and **ñ** with linear/quadratic kernels *v*_*i*_/*v*_*ij*_ and 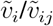. This is the argument submitted to Ψ in expression 2. If we approximate Ψ to first-order (linear), we know from previous work (Neri, 2010b) that:

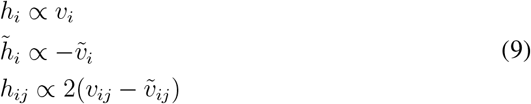

where ∝ stands for the same proportionality factor across the three expressions. These results prove reasonably robust for substantial deviations of Ψ from its first-order approximation, with the following caveats in mind. First, if 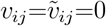 (i.e. Δ*R* is linear with no quadratic terms), the expressions above apply for any smooth Ψ, without the need for low-order approximations (Bussgang, 1952; Neri, 2010b). Second, readers may be tempted to infer that one may compute target-present and target-absent components of the second-order kernel separately and that, in similar fashion to the first-order kernels, these components may be simply expressed as ∝*v*_*ij*_ and 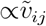. This is only true for a first-order approximation of Ψ, but quickly becomes inapplicable for small deviations from this low-order approximation (Neri, 2010b). For this reason, we only compute the composite second-order kernel *h*_*ij*_ derived from both target-present and target-absent noise fields.

We now proceed by rewriting Δ*R* for each rule in the form of equation 8. For the projection rule (equation 3):

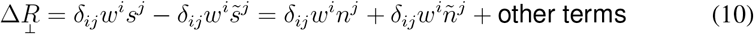

We therefore have:

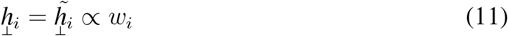

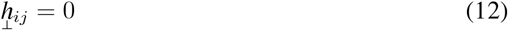

In words, the projection rule (linear-nonlinear detection mechanism) predicts indistinguishable first-order kernels, and a featureless second-order kernel. This is an established theoretical result (Ahumada, 2002). It has been observed in the laboratory under specific conditions (Beard and Ahumada, 1998; Neri et al., 1999; Neri, 2015) (see Section 3.2 and **Figure 2A**), however it is relatively uncommon that human empirical kernels comply with these specifications: it is more commonly found that 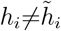 and *h*_*ij*_≠0 (Ahumada and Lovell, 1971; Dai et al., 1996; Barth et al., 1999; Abbey and Eckstein, 2002; Neri and Heeger, 2002; Solomon, 2002; Tjan and Nandy, 2006; Shub and Richards, 2009; Joosten and Neri, 2012; Joosten et al., 2016; Varnet and Lorenzi, 2022).

The absolute distance rule (expression 4) involves measure *D* (expression 1), which can be expanded as:

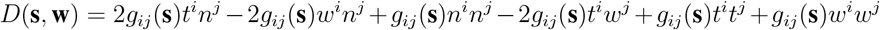

This expression may appear to conform to expression 8, however there is an important element of departure: *g*_*ij*_(**s**) depends on **s**, which in turn depends on its varying component **n**. To proceed further, it would be necessary to render this dependence explicit.

At the current stage in our knowledge and understanding of human perception, we do not possess an explicit description for the manner in which the metric varies across the perceptual manifold (see Section 5.4 for related considerations pertaining to the neural manifold). We may assume, for the purpose of attempting a minimal mathematical treatment, that there is an underlying connection with Riemannian characteristics, but we are not in a position to formulate an explicit specification for this connection. To derive expressions that provide any useful level of insight into the perceptual process, we must therefore adopt some approximations. The most important approximation we adopt here is the following:

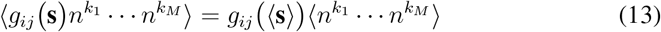

and similarly for the 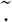 variants 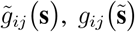, and 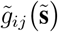, with *ñ*^*k*^ substituted for *n*^*k*^ when using 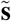. In words, it is assumed that the ambient metric is uncorrelated with local noisy modulations along any coordinate around ⟨**s**⟩ as measured with respect to **w**. To simplify calculations further, we also adopt the approximation:

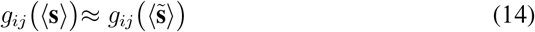

and similarly for 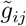. In words, it is assumed that the ambient metric around a given memory template (whether **w** or 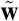) remains relatively stable regardless of whether it is queried with respect to the collective distribution of target-present stimuli, or with respect to the collective distribution of target-absent stimuli. This approximation is reasonable if the intensity of target signal **t** is relatively small compared with the noisy modulations applied to **s** and 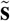: in the extreme case of **t**=0, 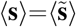, and equation 14 is trivially satisfied. The above approximation therefore only applies for low-to-moderate stimulus signal-to-noise ratios (SNR), which is the typical SNR regime adopted in kernel estimation experiments (Murray et al., 2002; Murray, 2011) (see Section 3.5 and **Figure 5F,I,J** for further analysis/discussion of SNR-related issues).

With the above approximations, the entire geometry of the perceptual manifold is summarized by only two metric specifications (⟨**s**⟩ and 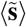 are fixed for a given configuration/characterization): *g*_*ij*_, which captures the ambient geometry around the memory template **w** associated with the perceptual representation of the target-to-be-detected, and 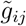, which captures the ambient geometry around the target-absent template 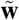. This degree of approximation may seem excessive, however we demonstrate that it is adequately matched to the degree of resolution afforded by current empirical measurements of relevant perceptual phenomena (Section 3). In the absence of explicit knowledge about the metric connection and its dependence on stimulus attributes, our only point of reference for empirical validation is provided by first-order and second-order kernel estimates. These estimates are coarse in the following sense: they cannot be obtained for single-shot presentations, but rather must be constructed on the basis of several input-output pairings across thousands of trials (Section 2.5). The resulting degree of characterization is compatible with the degree of approximation afforded by our analytical treatment here (see Sections 3.4, 4.1, and 5.1 for further discussion of relevant issues).

We can now approximate Δ*R* for the absolute distance rule as:

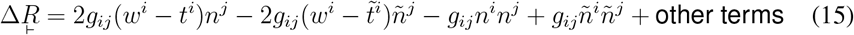

which conforms to equation 8. Notice that we only allow for this approximation to apply in the context of operating across many trials, which is how we calculate kernels from data (equations 6–7 in Section 2.5). Using equations 9, we have:

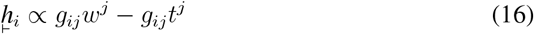

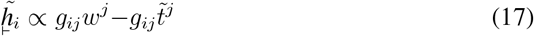

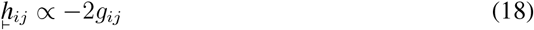

By adopting a similar procedure, we obtain the following approximation for the relative distance rule:

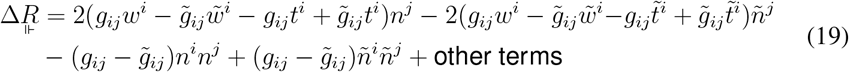

and the following kernel predictions (again from equations 9):

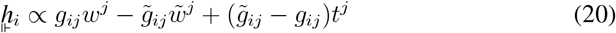

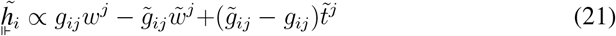

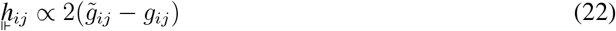

At this stage, we make the following observation: the relative distance rule subsumes the projection rule as a special case for 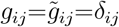 and 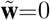. This can be easily verified by substituting those conditions into the expression for 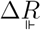, which returns 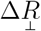 (10), or equivalently into the equations above for 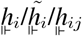, which return 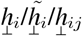 (equations 11–12). We can further relax the above conditions: **w** can be easily made to equal the zero vector by simply shifting the coordinate frame, making condition 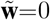 inconsequential; 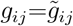 need not be equal to *δ*_*ij*_, as we can absorb *g*_*ij*_ into **w** by assigning to each element *w*^*i*^ the value returned by *g*_*ij*_*w*^*j*^ (in Euclidean space with Cartesian coordinates, vectors and covectors can be interchanged). We can summarize the above by stating that the relative distance rule collapses onto the projection rule when 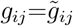. In the next section (2.7), we provide a geometrical interpretation of this fact.

The above observation does not apply to the absolute distance rule: 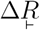 can only be made to equal 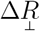 for the degenerate case in which they are both identically 0 (via setting *g*_*ij*_=0), which is not a viable choice of decision strategy (the system carries no discriminatory power).

### 2.7 Geometrical interpretation

We can visualize all three decisional rules within a common geometrical framework. We first imagine placing vectors **w**, 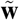, and **s** onto the coordinate frame (**Fig. 1B**). Without loss of generality, we can arbitrarily place the origin of the frame at 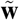, so that 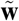 is the zero-coordinate vector. The projection rule amounts to projecting **s** onto **w** (black segment in **Fig. 1B**). The absolute distance rule amounts to measuring the (squared) distance between **s** and **w** (green segment in **Fig. 1B**). The relative distance rule amounts to measuring the distance between **s** and **w** (as in the absolute distance rule), then measuring the distance between **s** and 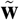 (red segment in **Fig. 1B**), and finally taking the difference between these two distances.

On the flat plane endowed with a Euclidean metric (bottom part of **Fig. 1B**), the relative distance rule is equivalent to the projection rule (observation at the end of the previous section): when 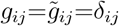 (Euclidean space), the two rules return identical Δ*R* values. In curved space (or a space endowed with curvilinear coordinates, see Section 5.4 for futher discussion of this point), the two rules differ greatly: in the example shown in **Fig. 1**, distance measurements in flat space (**Fig. 1B**) lead to the conclusion that **s** is more similar to 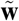 because it is closer to this vector (red segment is shorter than green segment); in curved space (**Fig. 1A**), distance measurements lead to the opposite conclusion that **s** is more similar to **w** because it is closer to this vector (green path is shorter than red path).

In our framework, the difference between flat space and curved space is captured by 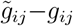: if this difference is 0, the components of the metric are flat (invariant across space) and the geometry collapses onto **Fig. 1B** (although see Section 3.4 for a demonstration of how this statement is not entirely accurate); if this difference is not zero, the components of the metric vary across space and the geometry lives on the curved surface in **Fig. 1A**. For the relative distance rule, the second-order kernel reflects precisely that difference (equation 22), thus making it a useful empirical descriptor of curvature in perceptual space. This observation is, in a nutshell, the primary motivation for the theoretical framework proposed here.

## 3 Experimental framework

### 3.1 Structural failures versus fitting exercises

Our goal is to construct a theory of perceptual processing, not a specific model of this phenomenon. To arbitrate between competing theories, we must avoid engaging in the class of fitting exercises that are often utilized for model selection, because we want mathematical structure, not parameterization, to bear on the arbitration process (Roberts and Pashler, 2000). This means that, if we exclude theory X, we do this because theory X is structurally incapable of accounting for the empirical evidence: it is the architecture of theory X that cannot capture the experimental result, not a specific choice of parameter specification for that architecture (Roberts and Pashler, 2000; Veksler et al., 2015).

For this type of selection programme to be enforceable, the most effective class of relevant experimental findings is such that a specific measurement is either 0 or not 0, and competing theories predict that it should be either 0 or not 0. When a theory predicts 0 (or not 0) and the empirical evidence returns not 0 (or 0), we speak of a “structural failure” or “architectural failure” of that theory (Neri, 2015). Notice that structural/architectural failures of this kind are not all equally compelling: if the theory predicts ≠0 and the evidence provides 0, this is weaker support for excluding the theory than the complementary scenario in which the theory predicts 0 and the evidence provides ≠0, because the absence of a measurable effect does not imply that the effect is absent.

If evidence of the kind outlined above is lacking, the only admissible evidence should be such that the theory makes relevant predictions that do not involve any free parameters (see Section 3.5).

### 3.2 Interdependence (and lack thereof) of first-order and second-order kernel estimates

We focus on two important empirical characteristics of kernel estimates: the presence/absence of potential differences between target-present and target-absent first-order kernels 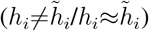, and the presence/absence of potential modulations within the second-order kernel 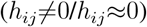. These quantities can be measured robustly with sufficiently large datasets (Ahumada and Lovell, 1971; Dai et al., 1996; Barth et al., 1999; Abbey and Eckstein, 2002; Neri and Heeger, 2002; Solomon, 2002; Tjan and Nandy, 2006; Shub and Richards, 2009; Joosten and Neri, 2012; Joosten et al., 2016; Neri, 2018b; Varnet and Lorenzi, 2022). By “absence” we mean not distinguishable from 0 within the resolution of our empirical measurements. Clearly, it is never possible to state that a given effect is completely absent: there always remains a residual possibility that the effect is actually present, but too small to be measured. Notwithstanding this inevitable difficulty with interpreting experimental results, we regard the resolution of our measurements to be sufficiently high for us to draw meaningful conclusions.

**Figure 2** presents three extreme cases of absence/presence for the characteristics just detailed, which encompass the entire range observed in the laboratory. The first two columns in **Figure 2** (panels **A,B,D,E**) come from the same visual experiment (Neri, 2018b), the only difference being stimulus contrast: high (8%) in **Figure 2A,D**, low (1%) in **Figure 2B,E**. Observers were asked to discriminate between target-present and target-absent stimuli structured as shown in **Figure 1C** (in compliance with the specifications of Section 2.2), both presented within the same trial (2AFC protocol), and briefly flashed (50 ms) in the near periphery (2–3 deg) to the left and to the right of a central fixation cross (please refer to the original publication (Neri, 2018b) for further details on stimulus specification). **Figure 2C** was computed from data collected during a virtually identical experiment (unpublished), with the only difference that no target was added to the target-present stimulus (on one out of three trials, unbeknownst to participants): **t**=0.

At high contrast, there is little to no measurable difference between target-present (blue) and target-absent (black) kernels (**Figure 2A**), and no measurable modulations within the second-order kernel (**Figure 2D**). At low contrast, there is a clear difference between target-present and target-absent kernels (**Figure 2B**), accompanied by substantial structure within the second-order kernel (**Figure 2E**).

The above transition 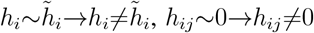 raises the following question: does 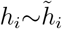 imply *h*_*ij*_=0, and/or does 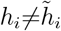 imply 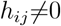?

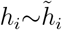 does not imply *h*_*ij*_∼0. It is easy to enforce 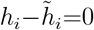: in the absence of a target signal, target-present and target-absent denominations remain undefined and 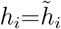 by definition, as also confirmed experimentally (**Figure 2C**). When analyzing experimental data under these conditions, trials must be randomly classified as target-present or target-absent, rendering this distinction only useful for gauging the extent of measurement noise (which is small, as indicated by nearly identical estimates in **Figure 2C**). The corresponding *h*_*ij*_ is nevertheless clearly structured (*h*_*ij*_≠0, **Figure 2F**).

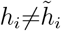 does imply *h*_*ij*_≠0 (at least empirically). It is never observed experimentally that 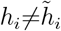 and *h*_*ij*_ ~ 0, except for pathologically high SNR values (Section 3.5, **Figure 5F,H**), when our approximations for deriving kernel estimates (see Sections 2.4 and 2.6) become inapplicable. In other words, if we classify all possible scenarios for the two characteristics under examination into binary absent/present categories, only three out of the four possible scenarios have been reported by previous studies: 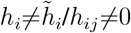 may be absent/absent (**Figure 2A/D**), present/present (**Figure 2B/E**), or absent/present (**Figure 2C/F**).

### 3.3 (In)compatibility of different rules with experimental evidence

The empirical results in **Figure 2** impose stringent constraints on candidate decisional rules, allowing us to arbitrate between them.

We start with the projection rule (⊥). This rule is incompatible with the presence of a measurable difference between target-present and target-absent first-order kernels: it dictates that 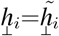 (equation 11). It is also incompatible with the presence of measurable modulations within the second-order kernel: 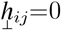(equation 12). Of the three experimental scenarios in **Figure 2**, it can only account for the high-contrast condition (left column). Furthermore, in its basic formulation, it is inconsistent with an iconic result in the perceptual literature known as the “dipper” effect (Appendix A). This rule must therefore be discarded because lacking sufficient flexibility to encompass the range exposed by laboratory experiments (Neri, 2015), despite being by far the most common model in sensory neuroscience (Heeger et al., 1996; Brunelli and Poggio, 1997; Dayan and Abbott, 2001; Ostojic and Brunel, 2011; Neri, 2015).

The absolute distance rule (⊢) allows for both absence and presence of a difference between target-present and target-absent first-order kernels: this difference is controlled by the 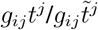 terms in equations 16–17. When *g*_*ij*_ is set to 0, 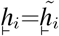; when *g*_*ij*_ is structured, 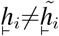. However, setting *g* _*ij*_ to 0 is not a viable option for three reasons. First, *g*_*ij*_=0 enforces 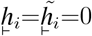, which is again contrary to experimental evidence: when 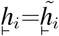, they are nevertheless structured (**Figure 2A,C**). Second, *g*_*ij*_=0 enforces 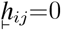, which is contrary to experimental evidence: the second-order kernel is often structured (**Figure 2E**), even when 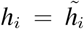(**Figure 2F**). Third, setting *g*_*ij*_=0 enforces 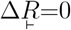 (equation 15), making the system unable to produce any meaningful measurement. Like the projection rule, the absolute distance rule must therefore be discarded because lacking sufficient flexibility.

In the relative distance rule (⊩), the difference between target-present and target-absent first-order kernels is controlled by the 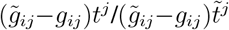 terms in equations 20–21. Furthermore, the first factor in these terms 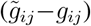 is proportional to the second-order kernel (equation 22). The relative distance rule therefore allows for both absence and presence of a difference between first-order kernels, and for absence and presence of structured modulations within the second-order kernel: when 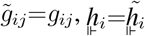 and 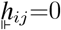; when 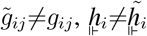 and 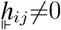. These two scenarios correspond to left and middle columns in **Figure 2**, respectively. The relative distance rule is also able to capture the scenario in the right column: when *t*^*j*^=0 (no target signal), 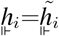 (capturing the experimental observations in **Figure 2C**) and 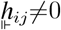 (capturing the experimental observations in **Figure 2F**), the latter provided that 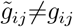. Finally, the relative distance rule cannot produce a scenario in which 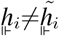 and 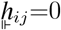, because the latter condition requires that 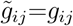, which in turn enforces 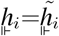. In summary, the relative distance rule is sufficiently expressive to capture all known experimental effects (**Figure 2**), and insufficiently expressive to capture effects that have not been reported experimentally (it also naturally accounts for the dipper effect, see Appendix A, and for error-producing kernels in the presence of twinned-noise, see Appendix B). In this sense, it is the most parsimonious metric rule that is consistent with empirical observations.

### 3.4 Computer simulations

To confirm and further study the geometrical framework outlined in **Figure 1**, alongside its connection with the 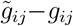 term in the kernel expressions for the relative distance rule (Section 2.7), we implemented this framework in software using Python. Input stimuli consisted of 13-element vectors specified as detailed in Section 2.2 and **Figure 1C–D**. Each stimulus was linearly projected to a 2-element vector via a pair of front-end filters similar to those in **Figure 1E–F**. The 2-element vector mapped to a 2D surface that could take any pre-defined shape in the embedding 3D volume. To compute geodesic distances on the surface, we incorporated the pygeodesic library into our software pipeline. We then applied the relative distance rule to produce binary responses, and analyzed the resulting dataset using identical procedures to those adopted with human data (Section 2.5).

When this procedure is carried out for a flat surface (**Figure 3A**), the resulting first-order kernels overlap (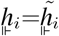, **Figure 3D**) and the second-order kernel is featureless (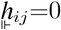, **Figure 3G**), resembling the experimental measurements in **Figure 2A,D**. We therefore refer to kernel measurements characterized by these features with the term “flat-looking”. When the same procedure is carried out for a curved surface (**Figure 3B**), the resulting first-order kernels differ (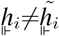, **Figure 3E**) and the second-order kernel is structured (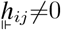, **Figure 3H)**, resembling the experimental measurements in **Figure 2B,E**. We refer to kernel measurements characterized by these features with the term “curved-looking”. When the target signal is removed, first-order kernels become identical (by definition) but the second-order kernel remains structured (not shown), thus capturing the experimental scenario in **Figure 2C,F**.

**Figure 3:**
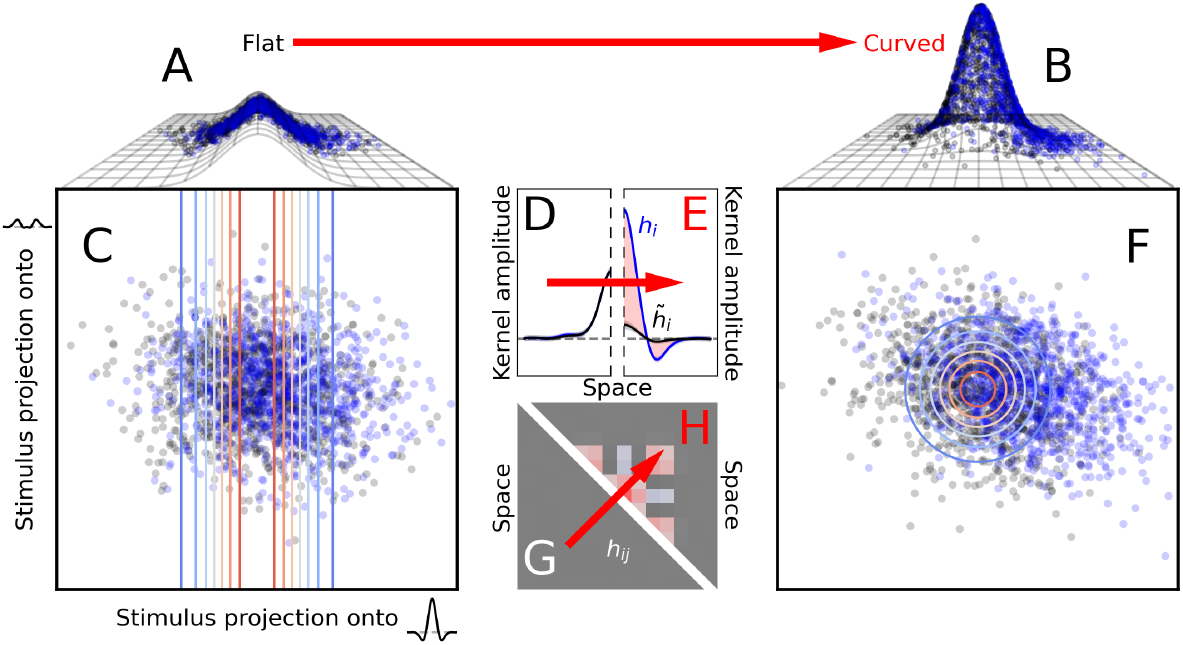
Simulated kernels for the relative distance rule capture the full range of human signatures. Stimuli were structured as in **Figure 1C**. They were projected onto the 2D plane in **C/F** (via linear projection onto the vectors illustrated by the small icons next to the axes in **C**), but distances were measured along geodesics on a potentially curved surface (**A/B**). Blue/black dots indicate target-present/target-absent projections. When the absolute distance rule is applied to distances measured on a flat surface (**A**), the resulting first-order (**D**) and second-order (**G**) kernels are “flat-looking,” resembling those obtained at high contrast in human experiments (**Figure 2A,D**). When the same procedure is carried out on a curved surface (**B**), the resulting kernels (**E,H**) are “curved-looking,” resembling those obtained at low contrast in humans (**Figure 2B,E**). **D,E** are plotted to the conventions of **Figure 2B**; **G,H** are plotted to the conventions of **Figure 2E**. Red arrows indicate the transition from flat (**A**) to curved (**B**) geometry.

In the above account of the empirically observed transition between left and middle columns in **Figure 2**, stimulus contrast is not explicitly modelled in any meaningful sense: this parameter is incorporated in the form of curvature, which is specified to be flat (**Figure 3A**) for the purpose of simulating high contrast, and curved (**Figure 3B**) for the purpose of simulating low contrast. In an alternative account of this experimental manipulation, it is possible to capture the resulting effect by specifying a single geometrical landscape with two hills (**Figure 4A**). In this account, contrast is explicitly modelled as a scaling factor applied to the stimulus projection: the low-contrast projection (red dots in **Figure 4B**) spans a smaller range than the high-contrast projection (black dots). Without modifying the geometry, the application of this scaling manipulation produces simulated kernels that adequately capture all salient characteristics of the experimental transition (**Figure 4C–F**).

**Figure 4:**
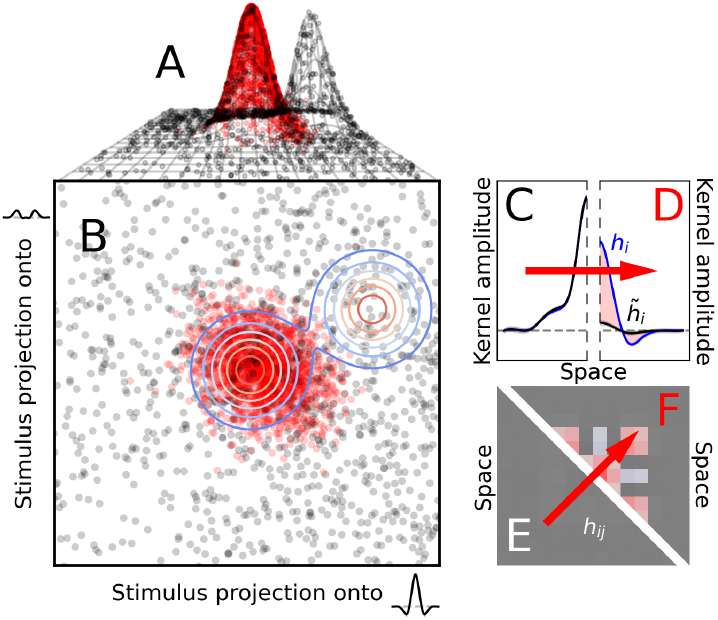
The transition from flat-looking to curved-looking kernels can be obtained without modifying the underlying geometrical landscape. In these simulations, the relative distance rule was applied to a surface with two hills (**A**). Low-contrast stimuli, indicated by red dots (without distinction between target-present and target-absent stimuli), project to the region occupied by only one of the two hills (**B**). The corresponding kernels (**D,F**) resemble those obtained in humans at low contrast (**Figure 2B,E**). High-contrast stimuli project to a wider region that spans both hills (black dots in **B**), producing kernels (**C,E**) that resemble those obtained in humans at high contrast (**Figure 2A,D**). While this kernel transition was produced by changing geometry from curved to flat in **Figure 3**, it is produced here without modifying the geometrical landscape, but rather as a consequence of sampling the geometry differently with low-vs high-contrast stimuli. **C,D** are plotted to the conventions of **Figure 2B**; **E,F** are plotted to the conventions of **Figure 2E**. Red arrows indicate the transition from the high-contrast (black dots in **A,B**) to the low-contrast (red dots) projection.

It may baffle readers that flat-looking kernels (**Figure 4C,E**) are produced by a curved geometry (**Figure 4A**). To understand this result, we must consider more closely the meaning of 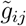 and *g*_*ij*_. In our theorical framework, flat-looking kernels are obtained when 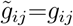 (Sections 2.7 and 3.3), where *g*_*ij*_ and 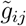 are ambient metrics (Sections 2.3,2.6). Templates **w** and 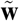 point to the summits of the two hills in **Figure 4A**. If we imagine placing the perceptual system at the summit of the hill sampled by low-contrast stimuli (red dots in **Figure 4A**), which corresponds to template 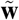, the system will experience ambient metric 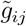 when tracing out geodesic paths to the surrounding stimuli. If we then move the system to the summit of the other hill, which corresponds to template **w**, the system will experience a very different ambient metric *g*_*ij*_ when tracing out geodesic paths to the low-contrast stimuli, because these paths will need to cross over to the hill corresponding to the 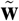 template. In other words, the collective projection for low-contrast stimuli looks very different when viewed from the top of the two hills. We can express this configuration as 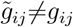, leading to curved-looking kernels (**Figure 4D,F**).

When contrast is raised, the collective stimulus projection (black dots in **Figure 4A**) becomes spread across the two hills and samples them to a comparable extent. When travelling from the top of one hill to the stimulus projection along geodesic paths, the system will therefore experience similar ambient metrics regardless of which hill it started from. In other words, the high-contrast projection establishes nearly symmetric sampling of the two hills. We can express this configuration as 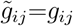, leading to flat-looking kernels (**Figure 4C,E**). This intuitive interpretation is consistent with our theoretical account (Section 3.3), and is confirmed by the simulations in **Figure 4** (we discuss this and related issues further in Sections 4.1 and 5.1).

The two scenarios simulated in **Figure 3** and **Figure 4** are conceptually different. In the first scenario, the transition from flat-looking to curved-looking kernels is effected by modifying the underlying geometry. In the second scenario, it is effected by modelling contrast explicitly, without modifying the underlying geometry. It is difficult to arbitrate conclusively between these two competing interpretations based on psychophysical evidence, because the manner in which the sensory process is interrogated by standard psychophysical procedures means that the underlying geometry cannot be reconstructed with certainty (see Section 5.1). In essence, when making binary choices of the kind considered here, observers sample only two regions within the perceptual manifold: the region around **w**, and the region around 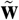. It is possible to attempt extended reconstructions provided some assumptions are incorporated into data analysis (Section 4), however it is not possible to conclusively pinpoint a specific architecture with certainty.

### 3.5 Non-parametric prediction of differential first-order kernel modulations from second-order kernel structure

Equations 20–22 show that our theory predicts a simple relationship between the two empirical characteristics that informed previous sections (difference between target-present and target-absent first-order kernels, structured modulations within second-order kernel). More specifically:

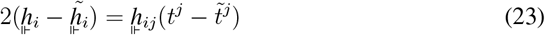

This relationship is not unique to the relative distance rule (Neri, 2010b) (an identical relationship applies to the absolute distance rule, see equations 16–18). Technically, it is also satisfied by the projection rule because, for this rule, both sides are equally zero (substitute equations 11–12 into 23). More generally, equation 23 applies to all models that are adequately captured by the analytical treatment adopted here Neri (2010b) (*ℋ* is captured by the Volterra formulation, Ψ can be approximated to first-order); in this sense, it speaks against models that are not accommodated by said framework, however it does not impose strong constraints for the exclusion of specific implementations, as it admits a relatively large class of models (but see Appendix C for examples of failures to comply with this equation). If confirmed by empirical data (see below), the significance of equation 23 therefore lies in its support for the applicability of the analytical treatment developed here to human sensory processing.

We know every term in equation 23: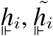, and 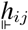 are estimated directly from data, while *t*^*j*^ and 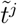 are specified by the experimenter (for the experiments considered here, 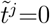). There are no free parameters involved.

This relationship holds very tightly in simulated scenarios: the shaded area (red tint) in **Figure 3E** shows the predicted difference between 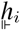 and 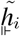 obtained by applying expression 23 to the second-order kernel in **Figure 3H**. A similar result is demonstrated in **Figure 4D**. In Appendix C, we present results from simulations involving circuit models, and show that sizable departures can occur under some simulated parameterizations.

When applied to real data, in general the prediction holds equally well: the redtinted region in **Figure 2B** shows an example for the low-contrast experiments discussed in Section 3.2, while **Figure 5A,B** show two examples from auditory experiments involving detection of loudness increments and decrements, respectively. **Figure 5E,F** show examples from unpublished low-contrast visual experiments similar to those in **Figure 2B**. Provided stimulus SNR is kept within a reasonable range (SNR=2 in **Figure 5E**), the prediction is accurate (red-tinted area matches difference between blue and black traces). When SNR is increased to non-physiological levels (SNR=10 in **Figure 5F**), the prediction fails completely (red-tinted area does not match difference between blue and black traces).

**Figure 5:**
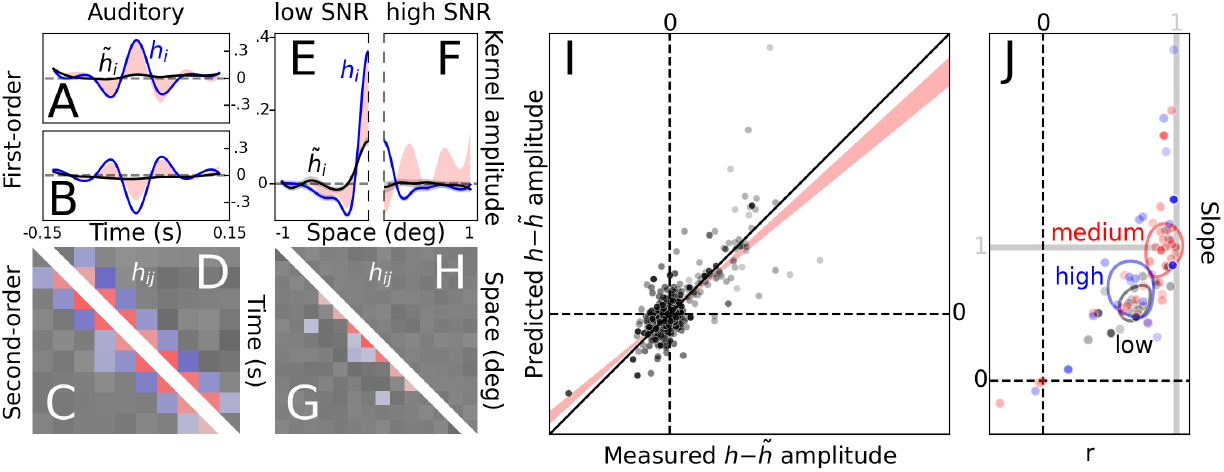
Second-order kernel maps directly to difference between target-present and target-absent first-order kernels with zero free parameters. First-order auditory kernels for detecting loudness increases/decreases (**A/B**) present clear differences between target-present (blue traces) and target-absent kernels (black traces). These differences are fully accounted for (red-tinted regions in **A/B**) by projecting the target signal onto the corresponding second-order kernels (**C/D**) as specified by equation 23. For a stimulus SNR value of 2, which is typical in psychophysical experiments (Murray et al., 2002), the match between measured and predicted difference is equally good in visual experiments (**E,G**; see also **Figure 2B**). When SNR is raised to a value of 10, which is well above typical laboratory ranges, the prediction fails (**F,H**). **I** plots point-by-point predicted (y axis) versus measured (x axis) differences between target-present and target-absent kernels from a wide range of experiments/conditions, obtained by aggregating ≈ 613K trials across four published studies (Neri, 2010c, 2013; Joosten et al., 2016; Neri, 2018b). The best linear fit (red-shaded area in **I** shows 95% confidence intervals around fit) is obtained for a slope of 0.86 and intercept ~ 0, with a correlation coefficient of 0.71. **J** plots slope (y axis) and correlation coefficient (x axis) separately for different kernels (one data point per kernel), sorted into three percentile clusters for low/medium/high SNR (low*<*2 ≤ medium ≤ 3*<*high). Oval contours reflect spread of different clusters (for visualization only). Optimal slope values of 1 (indicated by horizontal gray line) and correlation coefficient values approaching 1 (indicated by vertical gray line) are achieved for medium SNR levels (red cluster). Symbol opacity reflects square root of number of trials that contributed to its computation in **I,J** (low opacity for fewer trials). Kernels in **A,B,E,F** are plotted to the conventions of **Figure 2B**; those in **C,D,G,H** are plotted to the conventions of **Figure 2E**. Panels **A**–**D** were computed from 91K trials of published data (Joosten et al., 2016) aggregated across 10 listeners. Panels **E**–**H** were computed from 106K trials of unpublished data aggregated across 9 observers.

The above failure is expected, because our derivations assume low-to-medium-range SNR values (Section 2.6). Notice that SNR could be increased to uncharacteristically high values in the experiments of **Figure 5F** because they involved a very peculiar regime in which contrast was unusually low, remaining below detection threshold. In more typical conditions when stimulus visibility is not a limiting factor, SNR values as high as the one demonstrated in **Figure 5F** (SNR=10) would not permit kernel estimation, because the human observer would be nearly always correct on every trial. On the few trials on which the observer may respond incorrectly, this behavior would likely be attributable to stimulus-decoupled factors, such as blinking or inattention, that are not correlated with input noise. In this specific sense that pertains to viable threshold experiments, SNR values as high as 10 are not physiological (see also Murray et al. (2002)).

To gauge the applicability of expression 23 more quantitatively and across a wider range of experimental settings, we applied this prediction to *>*600K trials from various conditions/observers/experiments spanning four published studies (Neri, 2010c, 2013; Joosten et al., 2016; Neri, 2018b). **Figure 5I** plots predicted (y axis) versus measured (x axis) amplitude differences between target-present and target-absent first-order kernels. Points scatter around and along the unity line, indicating that the prediction generalizes adequately to a broad range of experimental configurations.

To further study the impact of SNR, we sorted this large dataset into three equally-sized percentile groups corresponding to different SNR levels (low, medium, high), and measured the slope/correlation-coefficient associated with each kernel in each group when plotted separately in the manner of **Figure 5I**. At low and high SNR (black and blue data points in **Figure 5J**), both slope and correlation coefficient fall below the optimal target value of 1 (black/blue ovals fall below the horizontal gray line and to the left of the vertical gray line in **Figure 5J**). At medium SNR (red data points in **Figure 5J**), these two characteristics are very close to 1 (red oval falls near the point of intersection between horizontal and vertical gray lines in **Figure 5J**). This result is consistent with our theory (see below).

As already noted above, our theory does not apply at high SNR. More specifically, equation 23 relies on a linear approximation of Ψ (Section 2.6). This approximation is only reasonable for SNR values below ≈ 3 (Neri, 2010b). At higher SNR values, it becomes necessary to expand the approximation for Ψ to higher orders, however the resulting expressions involve ratios between expansion coefficients (Neri, 2010b) (e.g. between second-order and first-order), and in the best-case scenario these are not easily estimated from empirical measurements (in the worst-case scenario, depending on the nature of the experimental design, they cannot be estimated at all). Our goal is to remain with zero-free-parameter predictions such as equation 23 (see Section 3.1), so we do not attempt higher-order expansions here.

At low SNR, the amplitude of the target signal is small, weakening the distinction between target-present and target-absent kernel estimates. This configuration inevitably leads to highly unreliable measurements of their difference (if present). In the extreme case of SNR=0 (as in **Figure 2C**), it is meaningless to measure slope/correlation for the prediction from expression 23, because these two characteristics are bound to scatter around 0 (the prediction becomes equal to 0 across the board).

It may be argued that the above results merely reflect differences in data mass allocation to different SNR groups: it is possible that, by chance or by unwitting design, more trials were collected for experiments at medium SNR. Under this scenario, the medium-SNR group would be trivially expected to show higher correlation coefficients as a consequence of more reliable measurements. We can exclude this interpretation on the basis of two observations. First, the number of trials allocated to each kernel estimate is slightly larger for the low/high SNR groups (Q1–median–Q4: 5.1K–5.8K– 7K/5.2K–5.8K–6.9K) compared with the medium SNR group (5.1K–5.6K–6.5K). Second, there is no obvious relationship between symbol opacity (which scales with square root of number of trials allocated to the corresponding data point) and vicinity to target slope/correlation-coefficient values across data points in **Figure 5J**. We therefore conclude that the trend exposed by the scatter plots in **Figure 5I,J** does not reflect artefactual differences in data mass allocation across conditions.

## 4 Reconstruction of underlying geometry directly from data

If we adopt some simplifying assumptions, we can reconstruct and visualize the manner in which the metric varies across the perceptual manifold directly from data, without introducing any free parameters. For this purpose, we therefore allow *g*_*ij*_(**s**) and 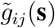 to vary with **s**.

### 4.1 Reconstruction in theory

Our first assumption is that the metric is rank 2, and that the eigenvectors are fixed for a given set of experimental conditions. For example, for the experiments involving contrast manipulations (left/middle columns in **Figure 2**), we assume that the same two eigenvectors apply to all stimuli across all contrast levels. To avoid excessive subscripting/superscripting in writing out explicit expressions for the projection of **s**, 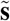, **w**, and 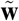 onto the two eigenvectors, we adopt the following conventions: the two scalar values obtained by projecting stimulus **s** onto the two eigenvectors are *s*_1_ and *s*_2_; the two scalar values obtained by projecting template 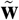 onto the two eigenvectors are *w*_1_ and *w*_2_; the two eigenvalues associated with *g*_*ij*_ are *λ*_1_ and *λ*_2_; similar notation applies to 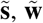, and 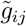 with the addition of the 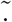 specifier.

Second, we assume that the perceptual process operates in accordance with the relative distance rule. Adopting the above assumption/notation, we can rewrite the output in response to stimulus **s** using the following simple expression:

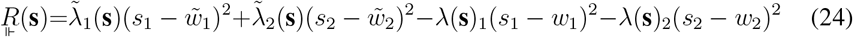

where the dependence on **s** has been displaced from 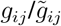 onto eigenvalues 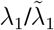 and 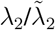, which essentially amounts to dimensionality reduction.

We express the Z-scored probability of producing a correct response as 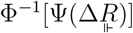 (Section 2.4) where Φ is the cumulative normal distribution function. We assume (as is customary) that Ψ is the cumulative Gaussian distribution function with standard deviation (SD) *α*, the SD of the internal noise source expressed as a multiple of the SD of the fluctuations induced onto the decision variable (*R*) by the external noise source (Green and Swets, 1966). We now approximate this function to first-order. For moderate SNR values, this approximation will be equivalent to a first-order approximation around 0 (which corresponds to SNR=0). The Z-scored probability will therefore be:

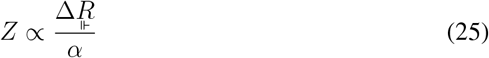

The proportionality constant above (∝) can be safely assumed to remain fixed across experimental manipulations that do not involve large changes in SNR (Neri, 2010b). We do not make this assumption for *α*: we allow here for the possibility that internal noise, as measured at the output of the decisional process, may vary across experimental manipulations. As explained below, this quantity can be estimated from data.

We now choose a specific target-present stimulus **s**_*⋆*_, and compute the Z-scored probability that it is classified as target-present when paired with all possible 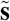:

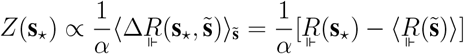

where the term 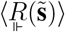 is clearly independent of **s**_*⋆*_.

Next, we choose a specific target-absent stimulus 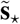 that projects to the same scalar values *s*_1_/*s*_2_ returned by the the projection of **s**_*⋆*_. Similarly to the procedure used with **s**_*⋆*_, we consider the probability that 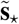 is classified as target-present when paired with all possible 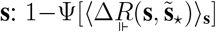. When Z-scored, this simplifies to:

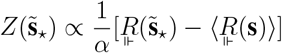

where the term 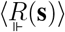 is clearly independent of 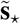.

We now average the two Z-scored probabilities above:

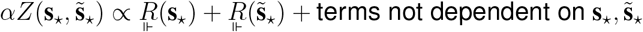

When written out explicitly (via equation 24), we obtain the following polynomial in *s*_1_/*s*_2_:

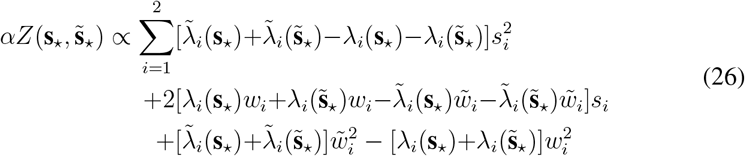

We want to study how the quadratic surface specified by *Z* varies as a function of small perturbations around *s*_1_/*s*_2_, to gain some insight into the manner in which 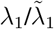 and 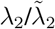 vary as a function of **s**_*⋆*_ and 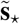. We make the reasonable assumption that the various *λ* factors vary more slowly with 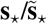 than the projected values associated with those stimuli (*s*_1_/*s*_2_). Failing this assumption, it becomes impossible to resolve/visualize *λ* variations, because they would exceed the resolution of our visualization strategy.

First, we examine the case in which the metric is constant across perceptual space (flat geometry): the *λ* factors do not depend on **s**_*⋆*_ and 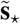, and 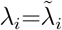. In this case, *Z* can be visualized as planar (equation 26 becomes linear in *s*_1_ and *s*_2_ because the leading (quadratic) term disappears). For the degenerate case 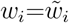, in which the system carries no discriminatory power (the two memory templates map to the same vector), *Z*=0 (chance performance).

Second, we examine the case in which the metric changes primarily with respect to the memory template from which distance is computed 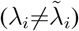, and less with respect to the specific choice of 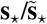 to which distance is computed 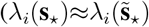 and 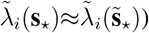. In this case, *Z* can be visualized as curved (equation 26 becomes quadratic in *s*_1_ and *s*_2_).

There is a third scenario in which the metric changes with respect to both memory templates and input stimuli. In general, *Z* will be visualized as curved under these conditions. However, it is possible for it to appear flat if **s**_*⋆*_ and 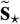 are chosen so that 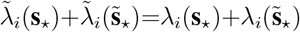. This is unlikely, but not impossible. Consider for example two zero-mean eigenvectors *u*_*i*_ (even) and *v*_*i*_ (odd). Choose **s**_*⋆*_ to be an indicator function for positive values of *u*_*i*_ (equal to 1 whenever *u*_*i*_ is positive, 0 otherwise), and 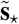 to be the negative of an indicator function for negative values of *u*_*i*_ (equal to −1 whenever *u*_*i*_ is negative, 0 otherwise). **s**_*⋆*_ and 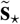 return the same values when projected onto *u*_*i*_ and *v*_*i*_, however they are sufficiently different from one another to allow for the reasonable possibility that 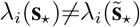. As long as 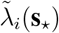 and 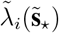 vary in mirror fashion to *λ*_*i*_(**s**_*⋆*_) and 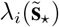 to keep their respective sums balanced, *Z* will look flat in the vicinity of the projected values *s*_1_ and *s*_2_. This scenario is not dissimilar from the scenario simulated in **Figure 4**, where a specific choice of stimulus distribution (high-contrast, black dots in **Figure 4B**) returns flat-looking kernels (**Figure 4C,E**) for a curved geometry (**Figure 4A**).

### 4.2 Reconstruction in practice

In practice, the reconstruction method must be subjected to an additional form of aggregation. Although in theory we can “choose 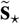 such that its projected values match those for **s**_*⋆*_,” this is not possible in practice: we cannot estimate the eigenvectors before running the experiment. In similar fashion, although in theory we can state “when paired with all possible 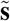,” we cannot repeat infinite measurements for a specific stimulus **s**_*⋆*_ when paired with all possible 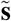, let alone do this for many choices of **s**_*⋆*_. When applying the reconstruction method to real data, we must bin the output range of *s*_1_/*s*_2_ and compute *Z* for all trials that fall within each bin. This practical implementation of the methodology further reduces its resolving power.

To estimate the eigenvectors associated with the metric, we rely on the assumption that they remain fixed throughout (for a given dataset). If we accept that the second-order kernel *h*_*ij*_ can be expressed as 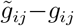, its eigenvectors will be those same eigen-vectors (except for the comprehensively flat case in which 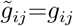 across the entire dataset). We can therefore estimate the two eigenvectors from the experimentally derived *h*_*ij*_ computed across the entire dataset (examples are shown by the two black traces near the axes at the bottom of **Figure 6A**). We can then project all stimuli used for a given experimental condition (e.g. low contrast) onto those eigenvectors, and compute *Z*.

**Figure 6:**
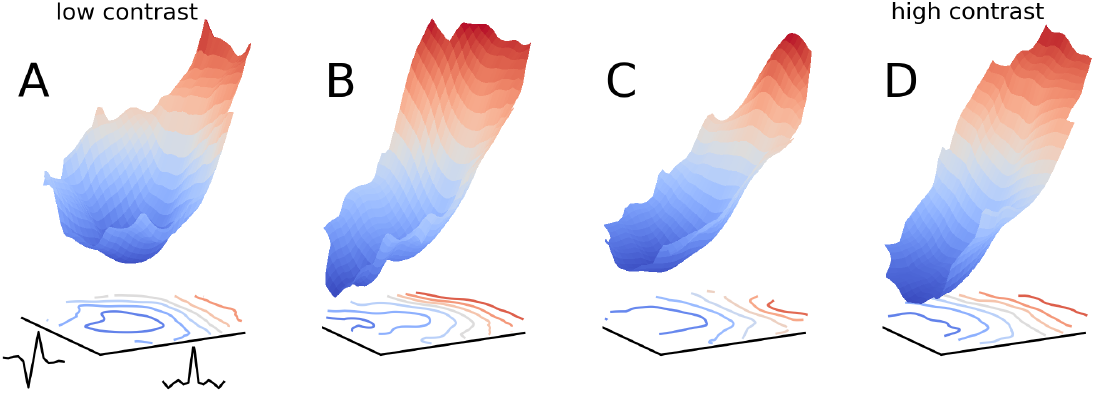
Reconstructed geometry for different contrast levels. Surface in **A** plots the Z-scored probability of classifying a given stimulus as target-present (z axis, corresponding to quantity *Z* in equation 26), scaled by internal noise *α*, as a function of the two values (mapped to x and y axes) associated with that stimulus when projected onto the two eigenvectors shown by black traces near x/y axes (these eigen-vectors were computed directly from the second-order kernel derived from the entire dataset across all contrast levels, see Section 4.2). In practice, projected values were binned for an arbitrarily chosen bin partition, and all stimuli/trials within a given bin contributed to the estimation of the corresponding probability (see main text for details). Consistent with corresponding kernel-based observations (left/middle columns in **Figure 2**), the reconstructed geometry at low contrast is curved (**A**), and becomes progressively planar/flat as contrast is increased (**B** to **D**). Surfaces were computed from 28K trials aggregated across 8 observers at SNR=1 (Neri, 2018b).

In equation 26, the lefthand side contains *αZ*, not just *Z*: to estimate the internal geometry of the response landscape (righthand side) prior to the application of the choice transducer Ψ, we must take into account (and factor out) internal decisional noise *α*. This quantity can be estimated experimentally using the double-pass technique (Burgess and Colborne, 1988; Neri, 2010a), which is the approach we adopt here (please refer to the original publications detailing internal noise characteristics for the datasets analyzed in this section, namely (Neri, 2018b) and (Neri, 2010c)). In the following, we therefore plot *αZ*.

**Figure 6** shows the outcome of applying the above procedure to the dataset already examined in Section 3.2, involving the transition from curved-looking to flat-looking kernels with increasing contrast (left/middle columns in **Figure 2**). In line with those observations, the reconstruction method returns a curved surface at low contrast (**Figure 6A**) that becomes progressively planar as contrast is raised (**Figure 6B–D**). Notice that, even when the geometry is predominantly curved (see blue-tinted region in **Figure 6A**), some portions of perceptual space may nevertheless be flat (red-tinted region).

In **Figure 7**, the reconstruction method is applied to a different dataset involving different levels of spatial uncertainty (Neri, 2010c). In this setting, kernel derivation is complicated by the fact that the target signal was not uniquely specified, in that it was allowed to vary its spatial location within a predetermined uncertainty window. In this case, kernel analysis can still be carried out using a technique termed signal-clamping (Tjan and Nandy, 2006), whereby noise samples are spatially registered to align with the target signal (Neri, 2010c). This ralignment procedure can only be applied to target-present noise samples, however, and not to target-absent samples, which may impact the applicability of the reconstruction process. Notwithstanding this limitation, it is interesting to observe that the reconstructed geometry varies from flat to curved with increasing uncertainty (**Figure 7A** to **Figure 7D**).

**Figure 7:**
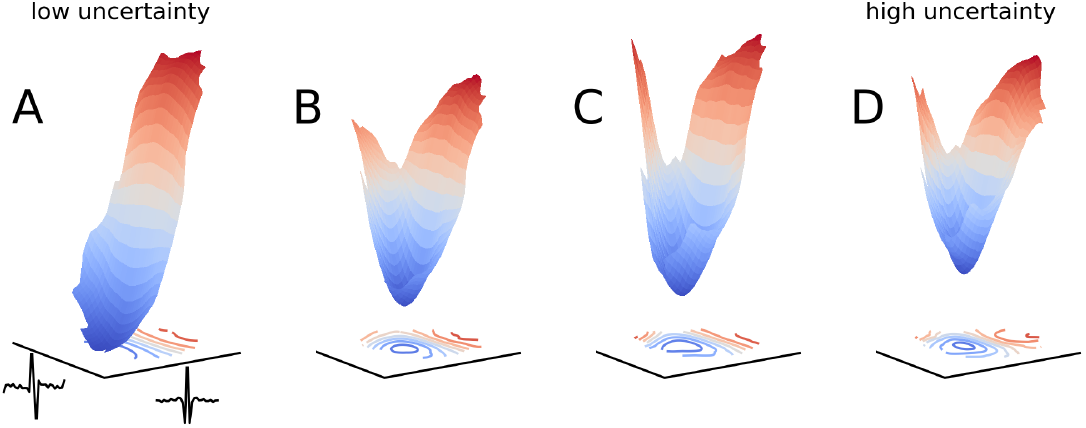
Reconstructed geometry for different uncertainty levels. Plotted to the same conventions of **Figure 6** using a different published dataset. When observers are allowed to focus on a restricted region of space where the target may appear (low uncertainty), the reconstructed geometry is flat (**A**). When the target is allowed to appear within increasingly wider regions of space (higher uncertainty), the underlying geometry becomes progressively more curved (**B** to **D**). Surfaces were computed from 94K trials aggregated across 10 observers (Neri, 2010c).

## 5 Discussion

### 5.1 Mathematics and perception

Perception in its most general sense (such as qualia) is far too complex a phenomenon to be available for mathematical treatment. It would already be an extraordinary achievement to establish a mathematical theory of perceptual processing in the sense of experimental psychology, that is in the sense of a phenomenon that can be quantified through behavior. If we restrict ourselves to the latter sense, we still find that not much progress has been made since the seminal work in the 1960s that led to the development of signal detection theory (Green and Swets, 1966) (SDT). A few other mathematical operators have been established for neuronal processing, such as energy models (Adelson and Bergen, 1991) and gain control (Carandini and Heeger, 2011), but their applicability to perceptual processing has been difficult to ascertain conclusively.

It seems relevant to ask: why has perception proven so difficult to capture using mathematical tools? There are various reasons potentially underlying this difficulty, ranging from the high variability associated with psychological phenomena to the inherent complexity and richness of perceptual experiences. At the same time, it should be recognized that these challenges have not hampered the mathematical characterization of neural activity to the same extent, despite those challenges being arguably equally relevant to neural representations: starting in the 1950s with the Hodgkin-Huxley model and subsequent developments in single-neuron biophysics, and progressing onto manifold accounts of multiunit activity in recent years (Kriegeskorte and Wei, 2021; Chung and Abbott, 2021; Langdon et al., 2023), it seems evident that neuromathematics (Ermentrout and Terman, 2010; Petitot, 2017) has provided compelling accounts of neurophysiological signals to a degree of detail and depth that has not been achieved for psychological and perceptual phenomena. The question is why.

We believe a large part of the answer lies in the extreme degree of information compression that is effected by animals when taking behavioral decisions (Zheng and Meister, 2024; Wu et al., 2016) (*ℋ* and Ψ in our formalism): animals receive a staggering amount of sensory data during natural behavior; this information is digested by their neural machinery to generate relatively rich perceptual representations; these representations are then used by animals to produce behavioral decisions in the form of motor outputs that ensure their survival, often involving little more than binary choices between discrete behaviors. The reduction in information rate associated with each of these transitions is so extreme that it can hardly be put into perspective. To provide a rough idea of the compression incurred by the translation from neural representation to behavioral output (second transition above), this can be estimated to involve a contraction from *>*1 gigabits/s to 10 bits/s (Zheng and Meister, 2024), which is at least a 10^8^-fold reduction in information rate (but see Sauerbrei and Pruszynski (2025) for a related discussion that takes into account motor control processes).

It follows from the above considerations that behavior provides an extremely impoverished readout of perceptual processes, thus making it difficult to impose serious constraints on candidate mathematical models (even low-dimensional ones) aimed at capturing behavior. For example, a sensitivity-based characteristic like the dipper function (Appendix A, **Figure 9B**) can be explained using a variety of different modeling schemes (Solomon, 2009), without clear evidence in favour of one or the other: the behavioral signature is just too poor a constraint to support effective model selection. In this study, we rely primarily on higher-dimensional behavioral descriptors (perceptual kernels, Section 2.5). These descriptors arguably provide a richer characterization of sensory behavior than sensitivity-based measurements (or at least complementary to the latter (Neri, 2018a)), however they too present important limitations: because of the compression associated with binary behavioral outputs, it becomes necessary to acquire many instances of such outputs to achieve a sufficient depth of empirical characterization (the kernels presented in this study come from dozens of thousands of trials).

The above considerations allow us to appreciate the non-trivial nature of the result in **Figure 5I-J**. This is a zero-free-parameter prediction of rich behavioral descriptors based on an articulated mathematical framework. It is remarkable that human behavior can be modelled to such an extent of precision using mathematics, and is in no way to be taken for granted: we have presented examples for which the theoretical prediction does fail under specific conditions (**Figures 5F**). Although this failure is to some extent expected from (and therefore indirectly consistent with) theory, it also demonstrates that the match between empirical descriptors and theoretical prediction is not a given (see also Appendix C): it is rooted in genuine compliance of human behavior with our theoretical treatment, which is ultimately based on mathematical expressions. If it is reasonable to find it unreasonable that mathematics is so profoundly effective at capturing natural phenomena in the physical sciences (Wigner, 1960), it seems reasonable to find it astounding that mathematics can capture so accurately a phenomenon as complex as human sensory behavior, notwithstanding the limited domain for which this result is demonstrated in the present study.

### 5.2 Relations to prior literature

It is legitimate to wonder whether/how the proposed framework may differ (or not) from existing accounts of sensory processing. More specifically, it may be argued that the geometrical account proposed in this study amounts to nothing more than rephrasing nonlinear frameworks for system characterization/identification (Golden et al., 2016; Marmarelis, 2004), such as the Volterra expansion. These frameworks have a long history in systems neuroscience (Marmarelis and Marmarelis, 1978; Westwick and Kearney, 2003), indeed the derivation of first-order/second-order kernels via noise-based reverse correlation techniques refers back to Volterra-like formulations (Section 2.6), both in physiology (Schetzen, 1980) and in psychophysics (Heath, 2000; Neri, 2004, 2010b).

Our approach in this study is fundamentally different for the following reasons. It was motivated by the specific goal of recasting perceptual processing using the simplest imaginable notion of what it means to take sensory measurements of the outside world: the notion of metric distance in perceptual space (Section 2.3). This is arguably the most natural choice for approaching the problem, to the extent that it may strike some readers as overly naive.

Once we commit to a unified account based on metric distance, the appropriate framework is intrinsic geometry. The Volterra expansion does not, *per se*, say anything about a specific notion of perceptual distance: it is a non-parametric general-purpose approximator for a wide class of functionals without any explicit connection with any specific construct, whether geometrical or otherwise. For example, the response of a system (*R* in our notation) may be successfully characterized by a Volterra expansion of the input stimulus (**s**) via the introduction of specific kernels (Marmarelis, 2004), but this does not say anything about the potential connection between the structure of those kernels and the perceptual operation that led to said structure.

To understand the above statement more concretely in direct connection with the material presented here, consider the transition from flat-looking to curved-looking kernels (left column to middle column in **Figure 2**). Both can be modelled using Volterra expansions, but this formulation does not say anything about whether their different characteristics originate from the same perceptual operation or from different operations, nor does it provide clearly interpretable information about what those operations may be. In our framework, the two characteristics originate from one and the same operation: measuring distances in perceptual space. The transition reflects a change in the intrinsic shape of perceptual space, but said space is subjected to the same operation for the purpose of generating a perceptual judgment.

For an additional example of the difference between our framework and the Volterra formalism, consider the manner in which we proceed when approximating the metric to low-rank in Section 4.1. Using this approximation/assumption, we can derive the eigenvectors of *g* directly from *h*_*ij*_. This approach is predicated on the notion that *h*_*ij*_ can be expressed as the sum of different *g*’s, however. Without an explicit connection between *h*_*ij*_ and *g*_*ij*_ (equation 23), there is nothing to tell us that we can apply the above procedure for inferring the eigenvectors of *g*_*ij*_ from *h*_*ij*_. This explicit connection is instantiated by our geometrical framework. In the Volterra formalism, the conceptual significance of the eigenvectors of *h*_*ij*_ remains unclear/unspecified.

More generally, our approach defines a simple geometrical framework that can potentially incorporate various cognitive phenomena (such as attention) in the form of intuitively graspable constructs, such as positional shifts of template vectors or disturbances in the distance-measuring process. Under a general-purpose approximator such as the Volterra expansion, those phenomena would merely involve compiling a list of induced kernel alterations with no unifying conceptual structure. At this stage, we do not know whether the potential of our framework for encompassing and understanding a wider range of cognitive phenomena may be realized or not, but we hope to address this issue in future work.

In a line of research that shares some conceptual characteristics with ours, some authors have proposed hyperbolic accounts of various perceptual phemonena, ranging from color vision (Yilmaz, 1962; Farup, 2014) to stereopsis (Luneburg, 1950; Indow, 1967). In these proposals, the perceptual metric is hyperbolic and specific cognitive phenomena, such as adaptation in color vision (Yilmaz, 1962) and movement speed in motion perception (Caelli et al., 1978), are associated with Lorentzian transformations. These accounts differ from ours in that, similar to special relativity, they live in a flat geometry: the associated second-order kernel is featureless, because the components of the metric are fixed. In our account, the observation of curved-looking kernels necessitates that said components vary across perceptual space. Furthermore, experimental evidence in support of those hyperbolic accounts is either not available or rather limited in its constraining power, at least in comparison with the depth of empirical scrutiny afforded by the methodologies employed in this study.

We conclude this section by drawing a clear distinction between the approach advocated here and existing mathematical treatments of various problems in vision science that rely on tools from differential geometry (Koenderink, 1984; Hoffman, 1989; Petitot, 2017). Those tools have played a central role in seminal work on the instrinsic geometrical structure of visual space (Koenderink, 1990). The frameworks developed by these prior studies “live” in the physical world, so to speak, in the sense of its ecological architecture (Gibson, 1979) and/or the optical image it projects (Koenderink, 1984; Turski, 2023), and as such they do not speak directly to the geometry of perceptual operations. Conversely, our framework lives in perceptual space: all relevant architectural constraints are imposed by the perceptual operation of computing geodesic distance, with little (at most indirect) input from the intrinsic geometrical structure of the physical world. Our account is also entirely local: the geometry is expanded around a single point in the optical image, with nothing to say about extended physical space and related phenomena (such as Gestalt rules of good continuation (Sarti et al., 2009)). In the future, it may be possible to specify the perceptual equivalent of field equations to describe how metric structure varies across space as a function of certain image characteristics (such as object boundaries or other salient features in the environment), however this seems a distant goal at the present moment. In conclusion, our framework is neither superior nor preferable to those discussed in this paragraph: it simply speaks to a different problem.

### 5.3 Relations to information geometry and statistical inference

In this section we highlight connections between our framework and recent developments in information geometry (Amari, 2016). Our treatment of perceptual processing must be formulated in statistical terms (Neri, 2010a), not only across trials (Sections 2.6, 3.4, 4.1, 5.1) but also with reference to any given trial (Section 2.4). In this sense, we can view perceptual decisions as measuring distances between distributions centred on 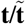 (with the stochastic component driven primarily by external noise) and 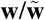 (with the stochastic component driven primarily by internal noise). This setup can be cast into a Riemannian framework with metric specified by Fisher information (Amari, 2016; Berardino et al., 2017; Ding et al., 2023), potentially providing some degree of specification for *g* in our formulation.

The above viewpoint is particularly productive in rendering explicit the connection between the geometry specified by the sensory representation of the input stimulus on the one hand (Sections 2.2–2.3), and the intrinsic variability associated with the decision process instantiated by the observer on the other hand (Section 2.4; see also Section 5.4 for additional considerations relating to the distinction between these two processes). Perceptual behavior depends on both factors, and their interplay can be understood via information geometry (Amari, 2016; Berardino et al., 2017). A rudimentary example of this connection was provided in Section 4, where the reconstructed perceptual surfaces were subjected to a scaling factor controlled by internal noise *α* (see also Seriès et al. (2009) and Berardino et al. (2017) for related approaches to Fisher information). Particularly if we imagine that *α* may vary across the surface, it becomes apparent how the curvature of perceptual space (as inferred from behavioral signatures in **Figures 6**–**7**) is impacted not only by what we regard as the intrinsic geometry of the sensory representation (curved landscape in **Figure 1A**), but also by the degree of noisiness associated with the manner in which the observer measures distances across the surface, and makes determinations about the presence/absence of specific target signals from such measurements (red/green paths in **Figure 1A**).

A related approach with a long history in visual neuroscience stems from ideal observer theory. When the memory template of the target signal is corrupted by Gaussian “memory noise” (Birdsall, 1960), the optimal strategy for statistical inference relies on a decisional variable that is a quadratic form of input vector **s** (equation 3.17 from Wilcox (1969), expressed here using notation consistent with the one adopted throughout this article):

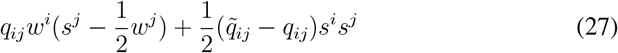

where *q*_*ij*_ and 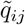 are the precision matrices associated with memory template **w** and with input noise, respectively. For Gaussian specification, these matrices are inverses of the covariance matrices associated with those processes (Wilcox, 1969), rendering obvious their link with Fisher information (Seriès et al., 2009). The connection with the framework developed here becomes apparent when we express 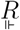 (equation 5) using the approximation *D*(*s, w*)=*g*_*ij*_*w*^*i*^*s*^*j*^ developed in Section 2.6, with 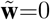 and 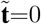 (detection task). We obtain that 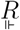 is twice expression 27 with *q* replaced by *g*. In other words, the framework developed here subsumes the case of an ideal observer with noisy memory (Birdsall, 1960; Wilcox, 1969), where the landscape specified by *g* reflects the statistical nature of memory templates and input processes along formulations that parallel those adopted in information geometry (Amari, 2016). In our framework, these different viewpoints are accommodated by one simple recipe for how perceptual processes operate when making decisions: they act like agents applying the relative distance rule to measured distances on arbitrary surfaces.

### 5.4 Curvature: intrinsic property, or curvilinear chart?

An important issue that was not explicitly addressed in this article relates to the coordinates adopted for specifying perceptual charts. Throughout this study, we have assumed Gaussian coordinates: those defined by orthogonal geodesics at a point. This assumption is implicitly incorporated into our simulations (Section 3.4) by measuring geodesic distance. Using this coordinate system, changes in the metric tensor transparently reflect genuine changes in curvature of the underlying geometry. What is the impact on our conclusions of allowing charts to be defined by curvilinear coordinates?

In a sense, the above issue is a non-issue, because it is not possible to distinguish between changes in curvature and changes in chart coordinates: we have no access to an underlying “true” geometry. In physics applications such as general relativity, for example, the notion of genuine curvature refers back to the physical world (matter distribution in particular). In psychological/perceptual space, there is no clear way of defining what is “real” and what is not: what is the difference between saying that the metric tensor defined in Gaussian coordinates varies across perceptual space because perceptual space is curved, and that the metric tensor defined in curvilinear coordinates varies across perceptual space despite said space being flat? In both cases, the perceptual system defines chart coordinates as well as underlying space, so from the viewpoint of metric measurements it is not possible to arbitrate between those two interpretations.

Notwithstanding the above considerations, it is possible to hypothesize a specific type of neural architecture for which the distinction between coordinate-induced and curvature-induced changes of the metric tensor takes on provisional meaning. If we posit the existence of an independent neural geometry at the level of brain activity, for example as represented by the spiking output of a neuronal subpopulation within a relevant sensory area (Sohn et al., 2019; Vyas et al., 2020; Langdon et al., 2023), and further posit that this geometry exists as a distinct phenomenon from the perceptual process that instantiates charts on it for the purpose of measuring distance and generating perceptual judgments, then we can ascribe curvature-induced changes to the former representation and coordinate-induced changes to the latter process. In other words, we are hypothesizing the existence of a “neural reality” that is independent of the perceptual process, and we ascribe genuine curvature (or lack thereof) to this “real” space. The perceptual process then becomes an observer of said reality: this observer may produce measurements that appear curved, even though the underlying neural reality is flat, merely as a consequence of laying out charts with curvilinear coordinates. We discuss this hypothesis further in Section 5.5.

### 5.5 The distinction between sensing, perceiving, and performing perceptual judgments

Our theory involves several intermediate constructs and computations preceding the final motor output produced by the system (Section 2). A relevant question is to what extent these “hidden” processes occur independently of whether they result in an overt motor output or not. For example, we may ask whether the geometrical landscape is instantiated by the brain of an anesthetized/asleep/unconscious individual (Chaudhuri et al., 2019; Gardner et al., 2022): if visual stimuli are flashed in front of his/her eyes, is the associated brain activity organized around a geometrical structure that retains the same characteristics elicited in the conscious state? Similarly, we may ask whether a specific chart is explicitly selected and represented by the sensory process of an individual who is not required to make perceptual judgments: is the mere act of perceiving the visual stimulus accompanied by the instantiation of metric components dictated by a spontaneously selected chart? Further, we may ask whether memory templates for specific stimuli are created spontaneously upon presentation of noiseless exemplars, whether they require explicit instructions/directions (e.g. “this is what your target looks like”), and/or whether they are only represented at the moment of making a perceptual judgment as to whether a given stimulus conforms to the template or not.

Needless to say, we currently do not have answers to these questions, so we can only speculate. We put forward a bold hypothesis in which we distinguish three components/stages: brain activity devoid of manifold structure (no atlas, **Figure 8A**); impromptu representation of manifold structure with Riemannian tools and addition/refinement of task-specific templates (**Figure 8B**); temporary choice and deployment of a specific chart (**Figure 8C**). We associate these three stages with three distinct states in which the human observer may be operating.

**Figure 8:**
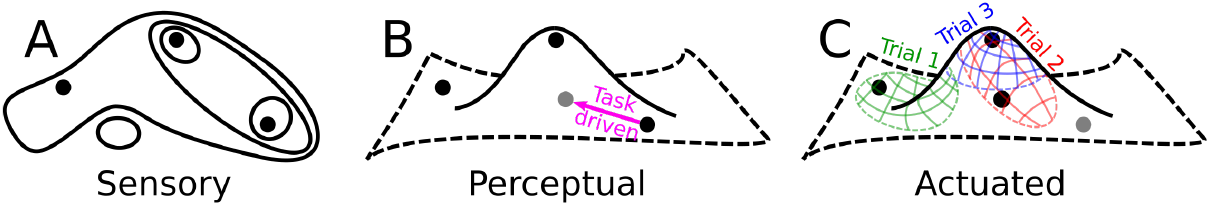
Speculative neurogeometrical scheme for the unfolding of perceptual processing from sensory encoding to motor action. We hypothesize that, in the absence of conscious perception, sensory representations are endowed with topological structure dictated by neural substrates (**A**). Perception enriches the topology with manifold architecture and a specific geometrical structure (**B**), which is encoded by stable descriptors in neural population activity. During task training, memory templates are recalibrated to represent task-relevant signals (magenta arrow). The geometry is not rendered explicit until the perceptual system is queried for the purpose of expressing a perceptual judgment, when it then selects a specific chart and performs actionable measurements on the underlying geometry (**C**). Every time the system is queried, a different chart may be selected (red/blue/green elements in **C**), which may appear in the form of trial-to-trial variations in neural activity. This scheme would account (among other things) for the co-existence of stable and heterogeneous characteristics at the level of neural activity (Low et al., 2018; Langdon and Engel, 2022).

We hypothesize that, in the absence of conscious (i.e. potentially actionable) perception, there is no manifold structure. This does not mean that there is no structure at all: neural activity may be organized according to an identifiable topology (**Figure 8A**) with relatable notions, such as conceptually “meaningful” subsets (e.g. the subset of all subparts belonging to a given object, the subset of all objects belonging to a given category, and so on). When we speak of manifold structure, we specifically refer to the representation of geometrical constructs that can be subjected to actionable computations. In the above, we use the term actionable with reference to a state/representation that can support action in the behavioral sense of motor output.

In the presence of conscious perception, geometrical structures are fully instantiated (**Figure 8B**), but they are not represented in computable coordinates with reference to a specific chart. This may be the state of an observer who is “looking at the world” without being assigned a specific behavioral task. Of course agents are constantly performing behavioral tasks, often self-initiated, but we hypothesize that the observer may be just “perceiving without acting” at least with reference to certain aspects of the sensory world. For example, the observer may be listening to a sound stream and performing actions in response to auditory information, however he/she may also be concomitantly sensing visual stimuli in the environment, without necessarily performing actions in response to visual information. We hypothesize that, under these conditions, neural activity is encoding the various geometrical constructs we have used in this study (such as the metric tensor) using representational schemes that are chart-independent.

Perception is inextricably informed by templates (sometimes termed priors (Mamassian et al., 2002)), some genetically encoded, others learnt throughout development. We hypothesize that these templates are spontaneously represented during conscious perception, however most of them will not be relevant to the specific behavioral task faced by the animal at a given time, so no distance measurements are computed for these templates. When observers are introduced to a new task, they will create new templates corresponding to the signals that are relevant to the task, possibly by modifying pre-existing templates (Hafting et al., 2005; Gardner et al., 2022) (magenta arrow in **Figure 8B**). For example, if observers are asked to detect a bright bar against a gray background, such as in the experiments considered in Section 3.2, observers will recall pre-existing templates for objects that are similar to a bright bar, clone one of them, and fine-tune its characteristics to align with the newly introduced target.

Finally, when observers are either prompted or forced to produce an explicit motor output based on a perceptual judgment, their perceptual system deploys a suitable chart (**Figure 8C**): all geometrical constructs within the charted region of perceptual space are assigned components specified by the selected coordinates, explicitly representing the machinery that is necessary to perform distance measurements. It then becomes possible to implement the operations described in Section 2, and generate corresponding motor behavior. The extent to which the selected chart aligns with geodesic paths may be controlled by cognitive factors, such as attention. For example, the curved-to-flat transition in **Figure 6** may reflect actual changes in the geometry of the neural representation, as it is driven by stimulus changes and is not under cognitive control (Neri, 2018b), while the flat-to-curved transition in **Figure 7** may reflect chart-induced misalignments associated with increasing uncertainty, as this factor is controlled by the state of the observer (Neri, 2010c) (see also Section 5.4 for a relevant discussion of related concepts from information geometry). Furthermore, different charts may be deployed on different instances when a behavioral output is produced in response to the same stimulus (different colors in **Figure 8C**), i.e. with reference to a stable geometry for the neural landscape. This chart-based interpretation would naturally explain the co-existence of stable and heterogeneous/transient characteristics at the level of neural population responses (Low et al., 2018; Langdon and Engel, 2022, 2025): the stable component would represent the underlying instrinsic geometry, while the transient and variable component would represent the deployment of ever-changing (but consistent when overlapping) charts.

At present, it is not possible to assess the applicability of the hypothetical scheme proposed above. However, geometrical techniques are gaining popularity in the analysis of neural population activity (Sohn et al., 2019; Vyas et al., 2020; Chung and Abbott, 2021; Genkin et al., 2025), potentially supporting characterizations that dovetail the behavioral framework developed here. In the future, we may have access to concomitant neural and behavioral measurements that are sufficiently rich to enable geometrical reconstructions such as those in **Figures 6**–**7**. Comprehensive measurements of this kind would support detailed assessment of the relationship between neural and behavioral components of sensory processing with reference to the hypotheses put forward in the preceding paragraphs.

## Acknowledgments

This study was supported by IIT (grant IVXX000701), the Agence nationale de la recherche (grants ANR-16-CE28-0016, ANR-19-CE28-0010-01, ANR-10-LABX-0087, ANR-10-IDEX-0001-02, ANR-22-CE93-0013), and CNRS.

## A The iconic “dipper” effect

### A.1 Theoretical considerations

Although our primary experimental point of reference is represented by kernel estimates (**Section 3.2**), we also consider one of the most iconic results in visual psychophysics: the dipper effect (Solomon, 2009). This is a counterintuitive phenomenon whereby the just noticeable difference *δ*_thr_ between a given stimulus of intensity *p* and another stimulus of intensity *p*+*δ* initially becomes gradually smaller as *p* increases, but then becomes gradually larger as *p* further increases beyond a certain value. In other words, *δ*_thr_ does not vary monotonically with *p*, as often assumed by coarse accounts of sensory discrimination such as Weber’s law, which dictates that *δ*_thr_ ∝ *p*.

We assume one fixed form of internal noise. More specifically we assume that, every time the perceptual system performs a distance measurement, this measurement is subject to an estimation error (Section A.2), analogous to taking measurements of physical quantities in the real world. In further analogy with many such physical measurements, we also assume that the magnitude of such errors scales linearly with distance to be measured: the larger the distance, the proportionally larger the standard deviation of the distribution associated with multiple measurements of said distance (Durgin et al., 2008). We refer to this type of noise with the term “Weber’s noise.”

We start with the projection rule. In the simplified case of a typical dipper experiment, all variables involved are simply scalars: 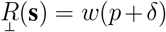 and 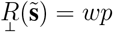, where *w* is the weight assigned by the system to the stimulus. To obtain approximate analytical expressions for *δ*_thr_, we recall that the resolving power of the system (its sensitivity) can be defined as follows (Green and Swets, 1966):

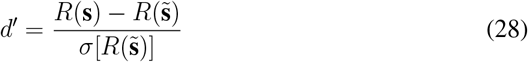

where *σ*[*x*] is the standard deviation of variable *x*. It should be noted that, technically, the above definition for *d*^*′*^ only makes sense when the distributions of variables *R*(**s**) and 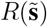 have comparable standard deviations (Green and Swets, 1966). This is not the case for Weber’s noise, but we adopt this definition here as a reasonable approximation, to be validated by computer simulations further below (**Figure 9**).

**Figure 9:**
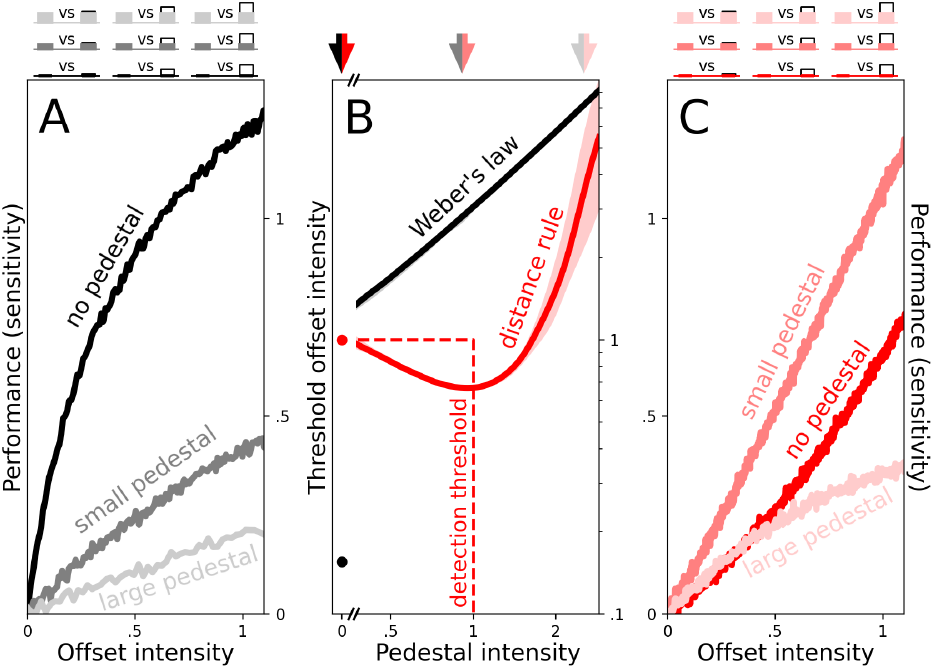
The relative distance rule naturally predicts the experimentally established dipper effect (Raab et al., 1963; Campbell and Kulikowski, 1966; Solomon, 2009). In this phenomenon, observers are asked to select the largest of two stimuli: one specified by a given pedestal intensity, the other one by an offset intensity added onto the same pedestal intensity (see icons above **A**, where offset is indicated by white rectangular element). As offset intensity is increased (x axis in **A**), discrimination performance (y axis) increases (traces in **A**). For a measuring system adopting the projection rule (black elements), and whose measurements are corrupted by intrinsic noise that scales with stimulus intensity (Weber’s noise), the rate of performance increase as a function of offset intensity scales with pedestal intensity: it is fastest for zero pedestal intensity (black trace in **A**), slower for small pedestal intensity (dark-gray trace), and much slower for large pedestal intensity (light-gray trace). The threshold offset intensity associated with a fixed performance level (y axis in **B**) scales linearly with pedestal intensity (x axis) in accordance with Weber’s law (black line in **B**). For a measuring system adopting the relative distance rule (red elements) and subjected to the same Weber’s noise (added to each distance measurement), the fastest rate of performance increase is not observed for zero pedestal intensity (red trace in **C**), but for an intermediate small value of pedestal intensity (dark-salmon trace in **C**). The corresponding threshold-offset-vs-pedestal-intensity function is therefore non-monotonic (red trace in **B**), with a trough that aligns with the pedestal intensity (dashed vertical line) that corresponds to the threshold offset intensity associated with zero pedestal intensity (dashed horizontal line, detection threshold). All numerical values are in units of this detection threshold. Shaded regions in **B** show 95% boundaries around median (line) across 100 iterations of simulated experiments.

Under the projection rule, 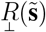 involves only one distance measurement (equation 3), so 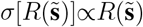 and 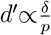. At threshold performance (constant *d*^*′*^), this implies *δ*_thr_ ∝ *p* (Weber’s law). In its most basic form, the projection rule cannot therefore account for the dipper effect.

We now consider the relative distance rule. Under this rule, 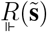 involves two distance measurements (equation 5): 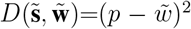 and 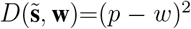. We therefore have that 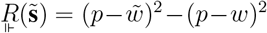 and 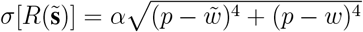, where we are taking into account both measurements in expressing *σ*, and *α* is a proportionality factor that reflects the overall intensity of internal noise (see Section 4.1). Without loss of generality, we can set 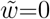. When substituting the above expressions alongside 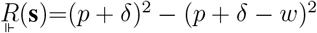 into equation 28 and setting *d*^*′*^=1 to enforce threshold, we obtain:

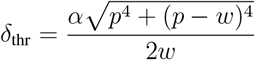

This expression is not monotonic: its derivative with respect to *p* vanishes for *p* = *w/*2. This value is similar to the detection threshold: *δ*_thr_=*wα/*2 for *p*=0. When *α* ≈ 1, which is typically the case in human sensory processing (Neri, 2010a), the dipper is therefore expected for a pedestal value near detection threshold, as reported experimentally (Raab et al., 1963; Campbell and Kulikowski, 1966; Solomon, 2009).

### A.2 Computer simulations

We confirmed the above conclusions with computer simulations. This is necessary because the expression for sensitivity (equation 28) does not necessarily reflect the process of measuring a psychometric curve in a 2-alternative-forced-choice pedestal experiment, and estimating *δ*_thr_ directly from said curve. Furthermore, the notion of internal noise that scales linearly with stimulus intensity creates difficulties around *p*=0 (detection): for the projection rule, for example, this leads to the anomalous result that the response to the pedestal becomes noiseless 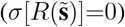. In computer simulations, we circumvented this issue by adding a small amount of “dark noise” (Barlow et al., 1957; Fuortes and Yeandle, 1964): a fixed factor added to the standard deviation of the internal noise source that remains present at zero stimulus intensity, and for which there is empirical support in human vision (Barlow et al., 1957; Srebro and Jaffe, 1968).

We simulated internal noise as exponentially distributed, with mean (and consequently standard deviation) set to stimulus intensity (*α*=1) plus the dark-noise factor mentioned above. Our choice of the exponential distribution was motivated by various considerations, for example the desirability of selecting a 1-parameter distribution with standard deviation that naturally scales with the mean. Furthermore, there is empirical evidence that internal noise in human sensory processing is Laplace-distributed (Neri, 2013), which is consistent with perceptual judgments based on differences of eponentially distributed variables. Finally, although very speculative and unspecified at this stage, it is not inconceivable that the process of geodesic exploration may share some distributional similarities with heat diffusion (Crane et al., 2017) and maximum-entropy energy allocation, making the Boltzmann distribution a potential candidate for modelling the spread of distance estimates over perceptual space. The current state of affairs is, unfortunately, that we do not know what the distribution of internal noise in human sensory processing conforms to. It has been assumed Gaussian without direct empirical evidence (Neri, 2013), however this assumption is almost certainly incorrect (Neri, 2013; Goris et al., 2014; Sanborn et al., 2025). For the purpose of our conclusions here, the choice of distribution is not critical, provided noise intensity scales with stimulus response.

We simulated end-to-end psychophysical experiments in which the proportion of correct responses was measured over 200 trials of scalar stimuli defined by *p* and *p*+*δ*. As offset intensity *δ* is increased (along x axis in **Figure 9A,C**), sensitivity (Z-scored proportion of correct responses, plotted on y axis) also increases (traces in **Figure 9A,C**). For a system that implements the projection rule (**Figure 9A**), the rate of increase becomes progressively slower as pedestal intensity *p* is increased (see Section A.1). For example, when *p*=0 (detection, see black icons above **Figure 9A**) sensitivity increases rapidly with offset intensity (black trace in **Figure 9A**). When *p* is increased to a slightly larger value (dark-gray icons), sensitivity increases more slowly with offset intensity (dark-gray trace), and this effect becomes more pronounced when *p* is increased to an even larger value (light-gray icons/trace).

From the above sensitivity-vs-offset traces, we measured threshold offset *δ*_thr_: the *δ* value for which sensitivity reaches a fixed arbitrary value (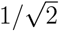in these simulations). When plotted (y axis in **Figure 9B**) against pedestal intensity *p* (x axis), this quantity increases linearly for the projection rule (black trace), in conformity with Weber’s law.

The relative distance rule presents a different behavior (**Figure 9C**). When pedestal intensity is increased from *p*=0 (red elements in **Figure 9C**) to a slightly larger value (dark-salmon elements), sensitivity increases more rapidly with offset intensity (compare red with dark-salmon traces). As pedestal intensity is increased further (light-salmon elements), sensitivity increases more slowly (compare red with light-salmon traces). This non-monotonic behavior is reflected by the red trace in **Figure 9B**.: as expected from theory (see Section A.1), this trace reaches a minimum for *p*=1, which corresponds to the value of *δ*_thr_ when *p*=0 (all values have been normalized by this specific *δ*_thr_ value).

## B Twinned noise

### B.1 Overview

Some prior studies have tested the applicability of different perceptual models to a “twinned-noise” variant of the classic signal-in-noise 2AFC paradigm, in which the two noise samples presented on any given trial are identical (Spiegel and Green, 1981; Swift and Smith, 1983; Burgess and Colborne, 1988; Eckstein et al., 1997; Ahumada and Beard, 1999; Beard and Ahumada, 1999; Johnson et al., 2002; Solomon, 2002; Burgess and Judy, 2003; Manahilov et al., 2005; Baker and Meese, 2012): **n**=**ñ**. The twinned-noise configuration makes specific predictions for the projection rule: if the observer were to conform to this decision scheme, there would be no coupling between input noise and perceptual decisions, because the modulation produced by input noise is subtracted out and 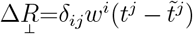 (no dependence on **n/ñ**). In other words, from the perspective of an observer that behaves according to the projection rule, it is as if the input stimuli contained no noise. Under this scenario, we expect improved performance (Spiegel and Green, 1981) and featureless kernels (Solomon, 2002) (see section B.2 for a detailed elaboration of the latter point).

Various studies have reported performance improvements (whether measured as increased sensitivity or decreased thresholds) when switching from “random-noise” (**n**≠**ñ**) to twinned-noise (Manahilov et al., 2005; Ahumada and Beard, 1999; Burgess and Colborne, 1988; Johnson et al., 2002; Burgess and Judy, 2003; Baker and Meese, 2012; Beard and Ahumada, 1999; Spiegel and Green, 1981; Eckstein et al., 1997), however other studies have failed to measure any reliable difference (Ahumada and Beard, 1997b,a; Pfafflin and Mathews, 1966; Watson et al., 1997), and some have reported poor replicability of previous results (Johnson et al., 2002; Burgess and Judy, 2003).

When taken collectively, the above-cited studies indicate that it is reasonable to expect some improvement in most experimental settings, however there may be specific configurations/protocols for which these improvements, if present, are not measurable. Even if we accept the general statement that performance improves with twinned-noise, this result is not overly useful for adjucating among competing perceptual models: in addition to the projection rule, many other rules predict improved performance for the twinned-noise configuration. For example, in our implementations of the absolute and relative distance rules in response to twinned-noise stimuli, the former can lead to improved performance under some parameterizations (although it can also lead to reduced performance under others), while the latter produces consistent (albeit small) improvements.

### B.2 Error-producing kernels from twinned-noise

A more powerful criterion for probing specific decisional rules is provided by first-order kernel predictions in the presence of twinned-noise. For the case of twinned-noise, kernels cannot be computed using the rules introduced in Section 2.5, because those rules are only meaningful in the presence of four types of noise average: ⟨*n*^*i*^⟩_✓_, ⟨*n*^*i*^⟩_*×*_, ⟨*ñ*^*i*^⟩ _*×*_, and ⟨*ñ*^*i*^⟩ _✓_. For twinned-noise, ⟨*n*^*i*^⟩ _✓_=⟨*ñ*^*i*^⟩ _✓_ and ⟨*n*^*i*^⟩ _*×*_=⟨*ñ*^*i*^ ⟩_*×*_, so that there are effectively only two types of noise samples: those associated with correct trials, and those associated with incorrect trials. We focus on the latter, because predictions for the former are essentially mirror images (with some scaling factor) of the latter. The closest equivalent of a first-order kernel for twinned-noise is simply given by ⟨*n*^*i*^⟩ _*×*_, the average of error-producing noise samples (Solomon, 2002). Below, we adopt the term “error-producing kernel” when referring to this descriptor.

As anticipated above, template-matching predicts featureless error-producing kernels (Solomon, 2002). In our formulation of the projection rule (which is equivalent to template-matching), this can be seen as a straightforward consequence of the fact that, under the twinned-noise configuration, 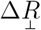 does not depend on input noise (see Section B.1). The “similarity” rule proposed by Watson and collaborators (Watson et al., 1997) predicts that error-producing kernels should be positively correlated with **t** (Solomon, 2002). In flat space, this rule is equivalent to the absolute distance rule (compare Figure 8 in (Watson et al., 1997) with **Figure 1B** here). In the presence of twinned-noise, Δ*R* for the absolute distance rule (equation 15) reduces to the following simple expression:

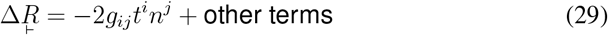

In flat space (*g*_*ij*_=*δ*_*ij*_), an incorrect response 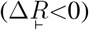 is therefore more likely to occur when the cross-correlation between **t** and **n** is positive (i.e. the first term in the above expression for 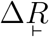 is negative), which aligns with the prediction for the similarity rule (Watson et al., 1997; Solomon, 2002). We confirmed this prediction with computer simulations (not shown).

In curved space, the sign of the term − 2*g*_*ij*_*t*^*i*^*n*^*j*^ depends not only on the cross-correlation between **t** and **n** but also on *g*_*ij*_, which in turn depends on the geometry of perceptual space and on the manner in which it is spanned by input stimuli (see Sections 2.7 and 3.4). We therefore predict that error-producing kernels may be either positively or negatively correlated with **t**, depending on how the two factors above are specified. We confirm and illustrate this prediction with computer simulations in **Figure 10A,C–D**. When input stimuli span a wide region of a curved surface (black dots in **Figure 10A,C**), the resulting error-producing kernel is positively correlated with the target signal (black trace in **Figure 10D**). When input stimuli span a smaller region of the geometry (orange dots in **Figure 10A,C**), the error-producing kernel becomes negatively correlated with the target signal (orange trace in **Figure 10D**).

**Figure 10:**
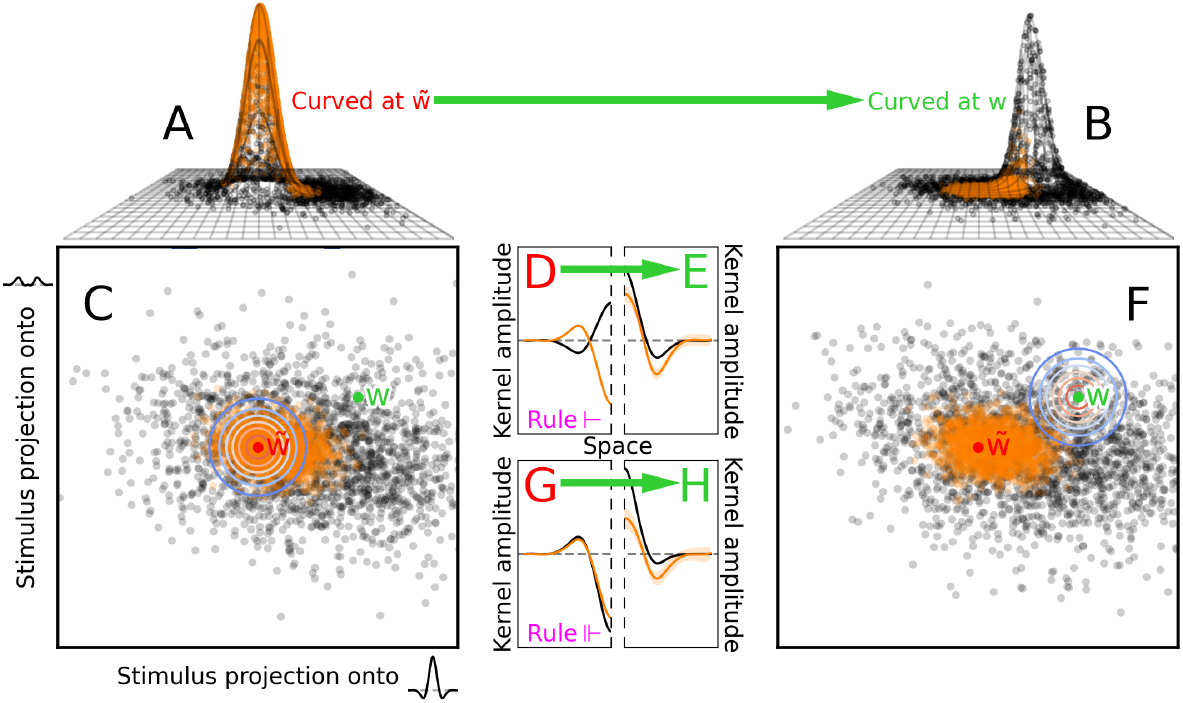
Error-producing kernels for twinned-noise. The simulations in **A,C** adopt the same parameterization used in **Figure 3B,F**, with the only difference that input noise was “twinned” (target-present and target-absent stimuli contained the same noise sample: **n**=**ñ**). Error-producing kernels for this configuration (computed by averaging noise samples associated with incorrect responses: ⟨*n*^*i*^⟩ _*×*_) are shown for the absolute (**D**) and for the relative (**G**) distance rules, when input stimuli span broad/restricted regions (black/orange dots in **A,B**, black/orange traces in **D,G**) of the landscape in **A**. Similar results are shown in **E,H** for a landscape (**B,F**) in which the curved region is shifted from the region around the projection of the target-absent memory template 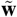 (indicated by red dot in **C**) to the region around the projection of the target-present memory template **w** (indicated by green dot in **F**). Human error-producing kernels (not shown) are negatively correlated with the target signal (Solomon, 2002), resembling the simulated kernels shown in **G. A,B** are plotted to the conventions of **Figure 3A**; **C,F** are plotted to the conventions of **Figure 3C**; **D,E,G,H** are plotted to the conventions of **Figure 3D** except for color specification (see above). Green arrows indicate the transition from high curvature at 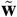 (**A**) to high curvature at **w** (**B**).

We can apply similar logic to the relative distance rule. In the presence of twinned noise, 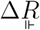 (equation 19) reduces to the following simple expression:

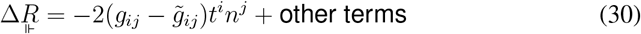

In flat space, this rule collapses onto the projection rule (Section 2.6), so it predicts featureless error-producing kernels (see above). This can be easily seen by setting 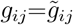 (the condition for flat space) in equation 30: the leading term becomes 0, thus removing any dependence on input noise.

In curved space, the term 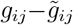 controls whether **n** should be positively or negatively correlated with **t** in order to increase the probability of an incorrect response 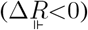. Generally speaking, if we consider noise samples for which the cross-correlation between **n** and **t** is negative, the term 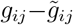 will invert the sign of the cross-correlation when curvature around memory template 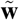 (captured by 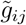) is larger than curvature around memory template **w** (captured by *g*_*ij*_). Under this scenario, the leading term in the expression for 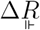 (equation 30) becomes negative, thus increasing the likelihood of an incorrect response. We therefore predict that error-producing kernels for the geometry outlined above should be negatively correlated with the target signal. We confirm and illustrate this prediction with computer simulations in **Figure 10A**, which depicts a geometry in which curvature around 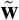 (marked by red dot in **Figure 10C**) is larger than curvature around **w** (marked by green dot in **Figure 10C**). Regardless of the extent spanned by input stimuli (black/orange dots in **Figure 10C**), the resulting error-producing kernels are negatively correlated with the target signal (black/orange traces in **Figure 10G**).

Based on the above logic, we expect that the sign of error-producing kernels for the relative distance rule should flip when curvature is distributed differently across the perceptual surface, such that curvature around **w** is larger than curvature around 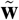. This geometry is depicted in **Figure 10B**. As predicted, the associated error-producing kernels are now positively correlated with the target signal (black/orange traces in **Figure 10H**). A similar result is obtained for the absolute distance rule (black/orange traces in **Figure 10E**).

### B.3 Comparison with human error-producing kernels

Solomon showed that, at least under the specific conditions of his experiments, the human error-producing kernel for twinned-noise is negatively correlated with the target signal (Solomon, 2002), leading this author to exclude the similarity rule (Watson et al., 1997) as a plausible model of the human process. We reach essentially the same conclusion here for the absolute distance rule with the caveat that, under some carefully chosen stimulus configurations and geometrical departures from flatness, this rule may generate human-like error-producing kernels (orange trace in **Figure 10D**). It is unclear how those configurations/geometries may translate to specific experimental tests of this prediction.

The relative distance rule aligns more effectively with human data: if we subscribe to the geometry underlying most of our treatments in previous sections (see **Figure 3B**), the associated error-producing kernels are consistently anti-correlated with the target signal (black/orange traces in **Figure 10G**). However, we have also shown that this rule can generate the opposite result when the geometrical landscape is manipulated in specific ways (black/orange traces in **Figure 10H**). It is unclear whether this should be regarded as a failure of the relative distance rule, or as potential for capturing human trends that have not so far been observed due to lack of relevant experimentation. To our knowledge, only one study (Solomon, 2002) has addressed this issue experimentally, and only for a very restricted set of stimulus parameterization (relatively high-contrast, small parafoveal probes). It is entirely possible that human error-producing kernels may show positive correlation with the target signal under different parameterizations/protocols. This remains an open question, which will hopefully be addressed by future investigations into the twinned-noise paradigm.

## C Simulated failures of 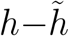 prediction

We have shown in Section 3.5 that human data largely conforms to the prediction from equation 23 (see **Figure 5**): for 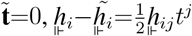. We demonstrate in this Section that it is possible to construct relatively simple and physiologically plausible circuit models that violate this prediction for mid-range SNR levels. We omitted/included specific components of the circuit in **Figure 11A** (see caption for description of individual components) to demonstrate their impact on simulated kernels. When the early nonlinearity (blue element in **Figure 11A**) and divisive normalization (red elements) are omitted, the prediction from equation 23 holds (**Figure 11F**). Kernels for this configuration are shown in **Figure 11B,D**. The addition of divisive normalization produces visible departure from this prediction (**Figure 11G**), particularly when SNR is raised from 2 to 3 (**Figure 11I**). Kernels for the latter configuration are shown in **Figure 11C,E**. When the post-filter static nonlinearity (orange element in **Figure 11A**) is swapped for the early static nonlinearilty (blue element), this manipulation also produces visible departures (**Figure 11H**).

**Figure 11:**
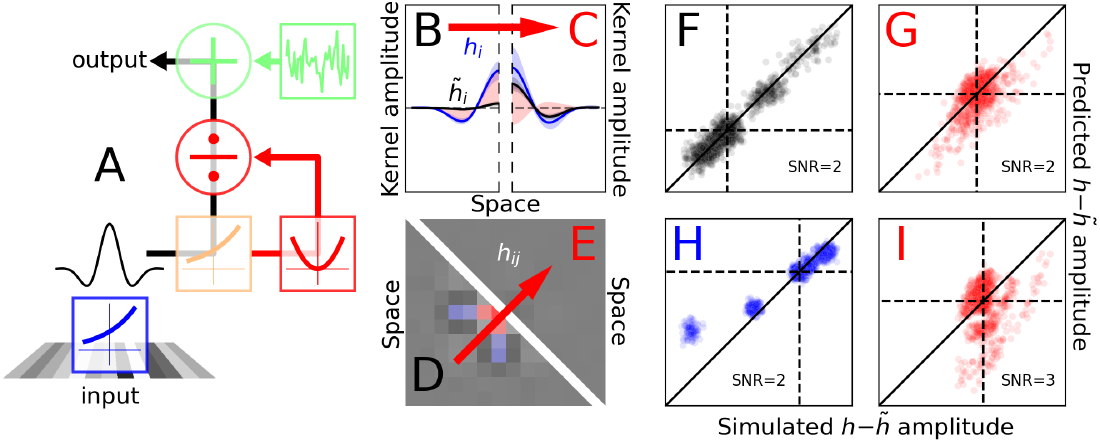
Compliance/departure with/from equation 23 for physiologically plausible circuit components. The simulated circuit model in **A** consists of the following components: early exponential (point) nonlinearity applied to the input stimulus (blue element); template-matching against Mexican-hat-shaped template (Gabor profile, normalized to RMS=1); scalar output from template-matching is subjected to further exponential nonlinearity (orange box); squaring and divisive normalization (red elements); late additive internal noise (green elements, *α*=0.5). Input stimuli were parameterized as described in Section 3.4 with SNR=2 (**F**–**H**) or SNR=3 (**I**); the circuit model was applied to both target-present and target-absent stimuli separately, and the stimulus associated with largest response was selected as target (Section 2.4). Divisive normalization of scalar value *x* was implemented as *x/*(*k*+*x*) with *k*=1 (**G**–**H**) or *k*=0.01 (**Figure 11I**), where *k* is expressed in units of SD of the output response to the target-absent stimulus (divisive normalization was omitted for the simulation in **F**). **F**–**I** plot results from 100 iterations of simulated experiments involving 10K trials each. **B,D** show kernels for simulations (**F**) excluding blue/red components in **A. C,E** show kernels for simulations (**I**) excluding only the blue component. **B,C** are plotted to the conventions of **Figure 2B**; **D,E** are plotted to the conventions of **Figure 2E**.

In general, departures of the kind demonstrated in **Figure 11G–I** are only observed provided SNR is at least in the mid-range between 2 and 3 (if SNR is reduced to 1, for example, the configurations adopted in **Figure 11** do not produce evident departures). Stimulus SNR values in the 2–3 range are not unusual in the psychophysical literature, so the simulations shown here are potentially relevant to laboratory settings. At the same time, we have never observed departures as large as those reported in **Figure 11I** when using mid-range SNRs (**Figure 5**), which indicates that the human process does not appear to implement, in general, the kind of nonlinearities that are exemplified by extreme configurations of the circuit model simulated here. We cannot of course exclude that such extreme scenarios may occur under experimental conditions that have not yet been tested in the laboratory.

## Notes

### Competing Interest Statement

The authors have declared no competing interest.

### Summary of Updates

Addition of a section to Discussion and two appendices

